# Polymyxin B lethality requires energy-dependent outer membrane disruption

**DOI:** 10.1101/2025.04.16.649083

**Authors:** Carolina Borrelli, Edward J. A. Douglas, Sophia M. A. Riley, Aikaterini Ellas Lemonidi, Gerald Larrouy-Maumus, Wen-Jung Lu, Boyan B. Bonev, Andrew M. Edwards, Bart W. Hoogenboom

## Abstract

Polymyxin antibiotics target lipopolysaccharide (LPS) in both membranes of the bacterial cell envelope, leading to bacterial killing through a mechanism that remains poorly understood. Here, we demonstrate that metabolic activity is essential for polymyxin lethality and leverage this insight to determine its mode of action. Polymyxin B (PmB) efficiently killed exponential phase *E. coli* but was unable to eliminate stationary phase cells unless a carbon source was available. Antibiotic lethality correlated with surface protrusions, LPS loss, and a significant reduction in outer membrane (OM) barrier function, processes that required LPS synthesis and transport. While the energy-dependent OM disruption was not directly lethal, it facilitated PmB access to the inner membrane (IM), which the antibiotic permeabilised in an energy-independent manner, leading to cell death. Finally, we show that the polymyxin resistance determinant MCR-1 prevents PmB-mediated LPS loss and OM protrusions and thereby renders the antibiotic ineffective.

## Introduction

The bactericidal activity of many antibiotics is dependent upon cellular metabolic activity [1,2,3,4,5]. For the efficacy of membrane-targeting antibiotics or peptides, however, it has been proposed that metabolic activity is not required, because intact membranes are presumed essential in all physiological states [6,7,8]. In support of this, many membrane-targeting compounds are effective against metabolically active and inactive bacteria [9,10,11,12,13].

Polymyxin antibiotics are the only membrane-targeting drugs used clinically against Gram-negative bacteria, but often lack efficacy *in vivo*, despite potent bactericidal activity *in vitro* [14,15,16,17,18]. Two polymyxins are used therapeutically, polymyxin B (PmB) and polymyxin E (colistin), both of which consist of a cationic cyclic peptide component coupled to an extracyclic peptide chain which in turn is linked to an acyl tail [14,15,18]. Both polymyxin antibiotics target LPS via high-affinity, long-lasting interactions, leading to disruption of the OM [18,19,20]. However, the nature of this OM disruption and the mechanism by which bacteria are killed by the antibiotic is largely unknown [14,15,18,21,22,23,24].

Several previous studies have shown that IM permeabilization, which is proposed to be required for killing, occurs after OM disruption [14,15,21,25,26,27,28]. Our previous work showed that LPS is the IM target of polymyxins, just as it is for the OM [27,28]. Whilst it is unknown how polymyxins cross the OM to access the periplasmic space, the predominant model is ‘self-promoted’ uptake, by which the cationic properties of the antibacterial disrupt cation bridges between LPS molecules, compromising barrier function and enabling antibiotic ingress through the OM [14,29,30].

Despite the membrane-damaging effect of polymyxins, they do not eradicate persister cells of *Escherichia coli* or *Acinetobacter baumannii* [31,32]. Stationary phase *E. coli* has also been shown to have a high level of polymyxin tolerance, suggesting that the bactericidal activity of these membrane-targeting antimicrobials is affected by metabolic activity and/or growth state [10]. Accordingly, metabolic activity was found to be required for colistin lethality at clinically relevant doses [13]. However, at concentrations above those typically achieved clinically (>10-50 µg ml^-1^), polymyxin lethality was found to be unaffected by growth phase or metabolic activity [11,32,33].

In summary, there are significant gaps in our understanding of how polymyxin antibiotics kill bacteria, and the requirement for metabolic activity remains unclear. We hypothesised that by resolving if and how metabolic activity was needed for polymyxin lethality, a clearer and more comprehensive understanding would emerge on the mechanism by which this class of antibiotics kill bacteria.

## Results

### Bacterial growth phase and nutrient availability affect PmB-mediated bacterial killing, membrane integrity and surface morphology

To assess the role of growth and metabolic activity in PmB-mediated killing, we took equal numbers of *E. coli* in exponential phase (high metabolic activity) or stationary phase (low metabolic activity) and exposed them to a clinically relevant concentration of the antibiotic (4 µg ml^-1^) in minimal medium only (MM) or with glucose (MM+G) to stimulate metabolic activity (Fig. 1A, Supplementary Fig. S1A) [34,35,36]. Exponential phase *E. coli* cells were rapidly killed by PmB regardless of whether glucose was present (Fig. 1A); and PmB efficiently killed stationary phase bacteria in the presence of glucose, albeit after a lag phase of 15 min during which there was no decrease in viability (Fig. 1A). Strikingly, however, there was no detectable killing of stationary phase *E. coli* during exposure to PmB in the absence of glucose, indicating that metabolic activity is required for the bactericidal activity of the antibiotic (Fig. 1A). This effect of glucose was dose-dependent, and the non-metabolisable glucose analogue 2-deoxyglucose did not support PmB-mediated killing of stationary phase *E. coli* (Supplementary Fig. S1B,C). Conversely, there does not appear to be a requirement for cell division for PmB killing, since MM without antibiotic (+/− glucose) did not support noticeable growth of *E. coli* from either exponential or stationary phase during the assay.

**Figure 1.**
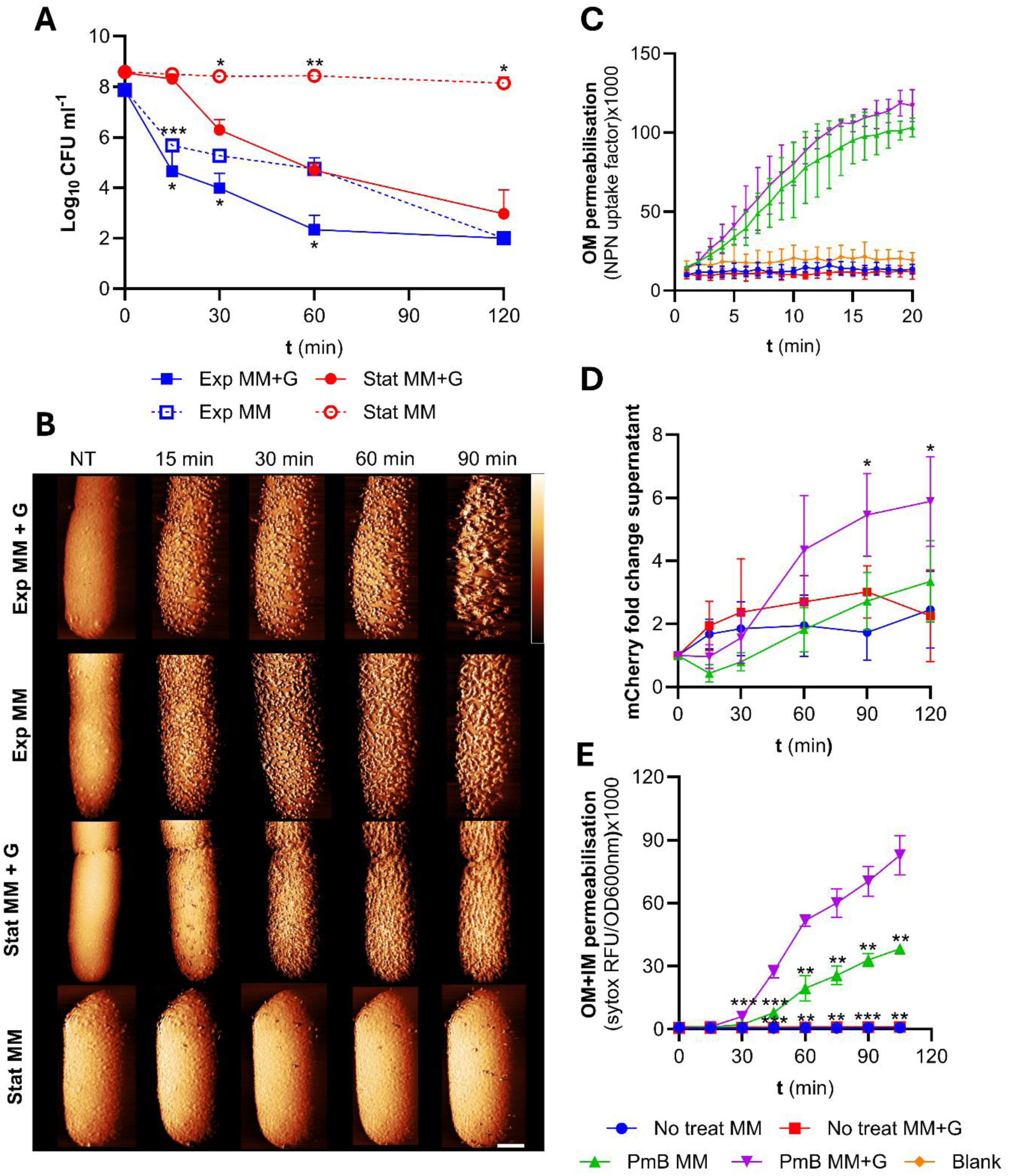
PmB lethality requires metabolic activity and is associated with significant morphological changes to the cell surface. **(A)** Survival of exponential or stationary phase *E. coli* exposed to 4 µg ml^-1^ PmB in MM +/− glucose, as determined by CFU counts. **(B)** AFM phase images showing exponential or stationary phase *E. coli* cells exposed to 2.5 µg ml^-1^ PmB in MM +/− glucose, shown as a function of time. Scalebar: 250 nm. Colour scale: 6 deg (row 1), 4 deg (row 2), 5 deg (row 3 and 4). Except for stationary phase cells without glucose, all bacteria were SYTOX positive by the end of the imaging (Supplementary Fig. S6A). Of note, a lower PmB concentration was used for AFM relative to other experiments, due to the low density of cells used in these assays. **(C)** OM lipid asymmetry/disorder of stationary phase *E. coli* cells during the first 20 min of exposure to 4 µg ml^-1^ PmB, as determined by uptake of NPN fluorescent dye. **(D)** OM disruption of *E. coli* _Peri_mCherry as determined by egress of the fluorescent protein mCherry into the culture supernatants of stationary phase bacteria exposed or not to 4 µg ml^-1^ PmB in MM +/− glucose. **(E)** Combined OM and IM disruption of stationary phase *E. coli* exposed to 4 µg ml^-1^ PmB in MM +/− glucose, as determined by uptake of the fluorescent dye SYTOX green. All experiments were replicated in n=3 independent assays. Error bars show the standard deviation of the mean. Significant differences were determined between stationary phase MM+G with PmB and each of the other conditions by two-way repeated measures ANOVA. P= *<0.05, **<0.01, ***<0.001, ****<0.0001, ns=not significant.

Using atomic force microscopy (AFM) to examine the effect of the antibiotic on the OM of living bacteria [37,38], we tracked the effects of the antibiotics in real time: PmB exposure of exponential phase *E. coli* resulted in a rapid and profound changes to the cell surface, with the appearance of numerous protrusions, regardless of the presence of glucose (Fig. 1B). By contrast, stationary phase *E. coli* only showed PmB-induced OM changes in the presence of glucose (Fig. 1B), correlating with the effects on bacterial viability (Fig. 1A,B). Scanning electron microscopy replicated the AFM observations, confirming these distinct, glucose-dependent phenotypes of stationary phase cells exposed to PmB, whilst the antibiotic caused surface protrusions in exponential phase *E. coli* regardless of the presence of glucose (Supplementary Fig. S2).

Next, using the small hydrophobic dye NPN [39] we showed that the binding of PmB to its lipid A target was unaffected by glucose or growth phase (Fig. 1C, Supplementary Fig. S3A). This finding was supported by mass spectrometry [27], which ruled out the presence of lipid A modifications in stationary phase cells that might reduce PmB binding, but intriguingly did show a large reduction in LPS abundance in cells exposed to PmB in MM+G (Supplementary Fig.S4). We further investigated changes to OM integrity by measuring the leakage of heterologously expressed mCherry (28 kDa) from the periplasm [40]. In the presence of PmB and glucose, stationary phase *E. coli* showed a significant increase of mCherry in the supernatant, compared with cells not exposed to the antibiotic (Fig. 1D).

Finally, we used SYTOX green to assess permeability of both the OM and IM during PmB exposure. As expected from the viability data (Fig. 1A), stationary phase *E. coli* showed significantly increased permeability in the presence of glucose, after a lag of 15 mins (Fig. 1E); permeability also ocurred in the absence of glucose, but at significantly lower levels than for cells exposed to the antibiotic in the presence of the sugar (Fig. 1E). Again, consistent with the viability data (Fig. 1A), PmB-induced IM permeability was unaffected by glucose in exponential phase *E. coli* (Supplementary Fig. S3B).

Taken together, these data show that PmB binds to *E. coli* regardless of metabolic state, leading to minor OM disruption sufficient to allow NPN ingress, but that bactericidal activity, OM protrusions and release of periplasmic protein require metabolic activity. Importantly, stationary phase viability and OM/IM permeation were similarly dependent on glucose across a diverse panel of laboratory and clinical isolates, indicative of a broadly conserved phenotype (Supplementary Fig. S5).

### PmB-mediated OM damage results in significant LPS loss

High-resolution AFM imaging of the OM of stationary phase *E. coli* showed a similar abundance and organisation of outer membrane protein (OMP) networks as previously reported [37,38] (Fig. 2A Supplementary Fig. S6B, note porous appearance of the OM in phase images). The porin networks were initially unaffected by PmB, but over time, the appearance of the PmB-exposed OM was dominated by protrusions at a scale >10 nm (Fig. 2A, Supplementary Fig. S6B,C). On these living *E. coli* cells, we did not observe any PmB-induced LPS rearrangement into hexagonal lattices, such as previously reported based on AFM on solid-supported, collapsed outer membrane vesicles [22,23,24].

**Figure 2.**
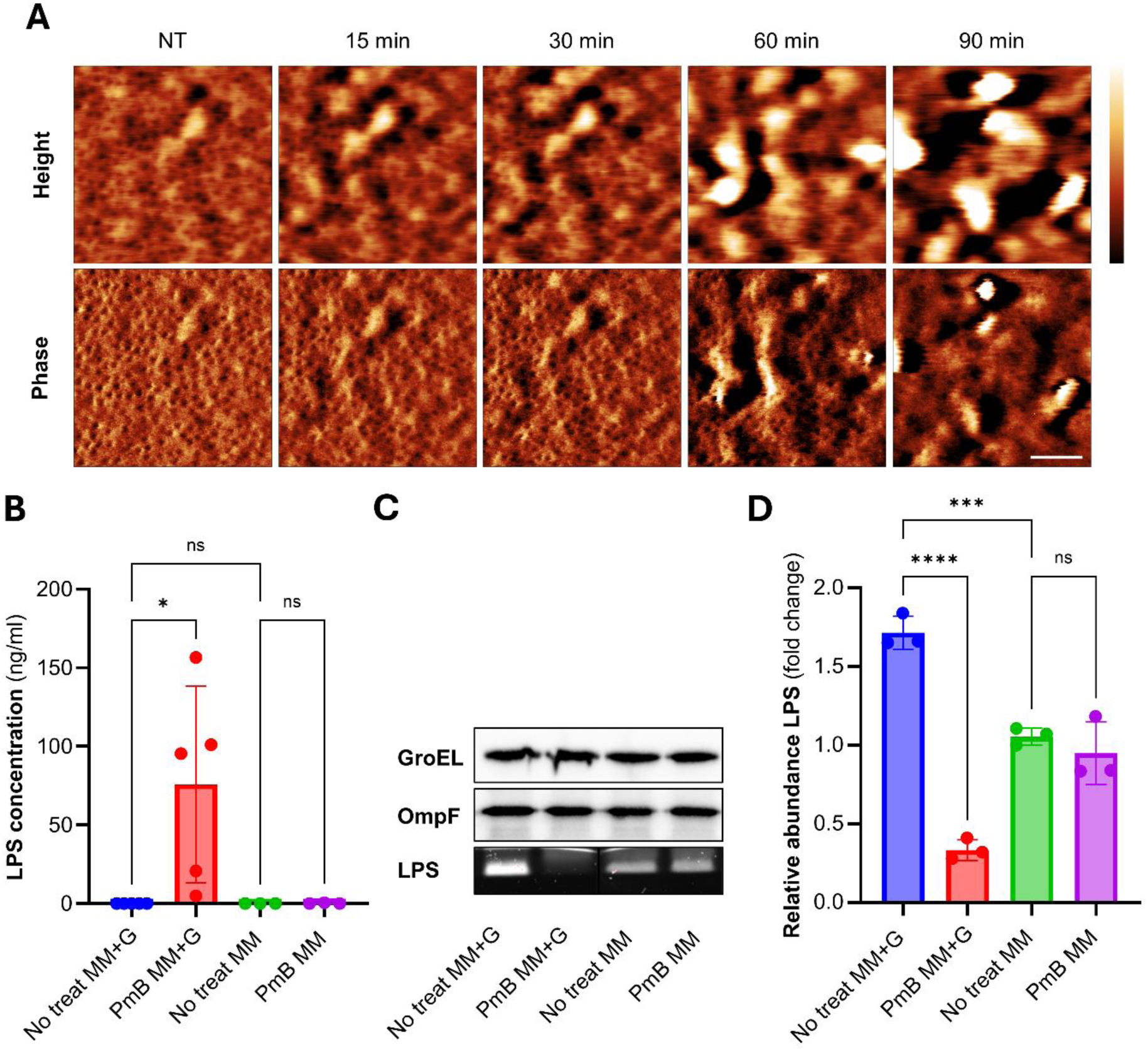
PmB-mediated OM disruption results in LPS loss without significant disruption of the porin network. **(A)** High-magnification AFM height and phase images of stationary phase *E. coli* exposed to 2.5 µg ml^-1^ PmB in MM+G, showing the porin network remains largely intact for at least 60 minutes of antibiotic challenge. **(B)** Kdo analysis of LPS recovered from filtered supernatants of stationary phase *E. coli* exposed, or not, to 4 µg ml^-1^ PmB in MM +/− glucose for 15 mins. **(C)** Levels of OmpF, GroEL and total LPS in *E. coli* exposed, or not, to 4 µg ml^-1^ PmB in MM +/− glucose for 15 mins. **(D)** quantification of LPS levels in *E. coli* exposed, or not, to 4 µg ml^-1^ PmB in MM +/− glucose for 15 mins. Scalebars: (A) 50 nm. Colour scales: (A) 3 nm (height) and 1 deg (phase). All experiments were replicated in n=3 independent assays. Error bars show the standard deviation of the mean. Statistical significance of differences was determined by one-way ANOVA, P= ***<0.001, ****<0.0001, ns=not significant.

Further examination of PmB-induced OM changes revealed that LPS was released from cells exposed to PmB in the presence of glucose, but not in the absence of the antibiotic or carbohydrate (Fig. 2B). Analysis of bacterial cells found a corresponding loss of LPS from *E. coli* exposed to PmB in the presence but not absence of glucose (Fig. 2C,D, Supplementary Fig S7), fully consistent with the mass spectrometry results reported above (Supplementary Fig. S4B). Surprisingly, the loss of LPS from *E. coli* was greatest at 15 min after PmB exposure and began to recover by 30 min, suggesting the bacteria attempt to repair the damage to the OM (Supplementary Fig. S8A). This coincided with the appearance of surface protrusions, which subsequently increased in abundance, up to IM permeabilisation as measured by SYTOX, (Supplementary Fig. S9). Moreover, LPS loss at 15 min was not due to cell lysis as, by contrast to LPS, there were no differences in levels of the cytoplasmic protein GroEL or OM porin OmpF across our experimental conditions (Fig. 2C).

Combined, these data revealed that PmB at least initially leaves the OMP networks intact but causes substantial and rapid LPS loss in the presence of glucose. PmB-induced LPS loss also occurred with *P. aeruginosa* and with exponential phase *E. coli* in MHB, indicative of a conserved process (Supplementary Fig. S8,S10).

### PmB-mediated LPS loss and bacterial killing require energy and LPS synthesis

The observed dependence on metabolic activity for PmB lethality suggests an energy-dependent bacterial response to PmB challenge. Since PmB-triggered LPS release required metabolic activity, we hypothesised that LPS synthesis and/or transport to the OM contributed to bacterial killing.

To establish energy dependence, we sought to distinguish between carbon– and energy-related effects. Using equimolar concentrations of various carbon sources in MM, we measured stationary phase *E. coli* survival under PmB challenge and, in parallel experiments without PmB, bacterial ATP production and overall LPS abundance (Fig. 3A). In these experiments, bacterial PmB susceptibility correlated with ATP production and with the amount of LPS associated with cells in the absence of PmB (Fig. 3A,B, Supplementary Fig. S11). In the presence of PmB, the amount of LPS dropped to lower levels in the presence of carbon sources that yielded higher ATP production (Fig. 3B, Supplementary Fig. S11), confirming the importance of energy for PmB-induced LPS loss and bacterial killing.

**Figure 3.**
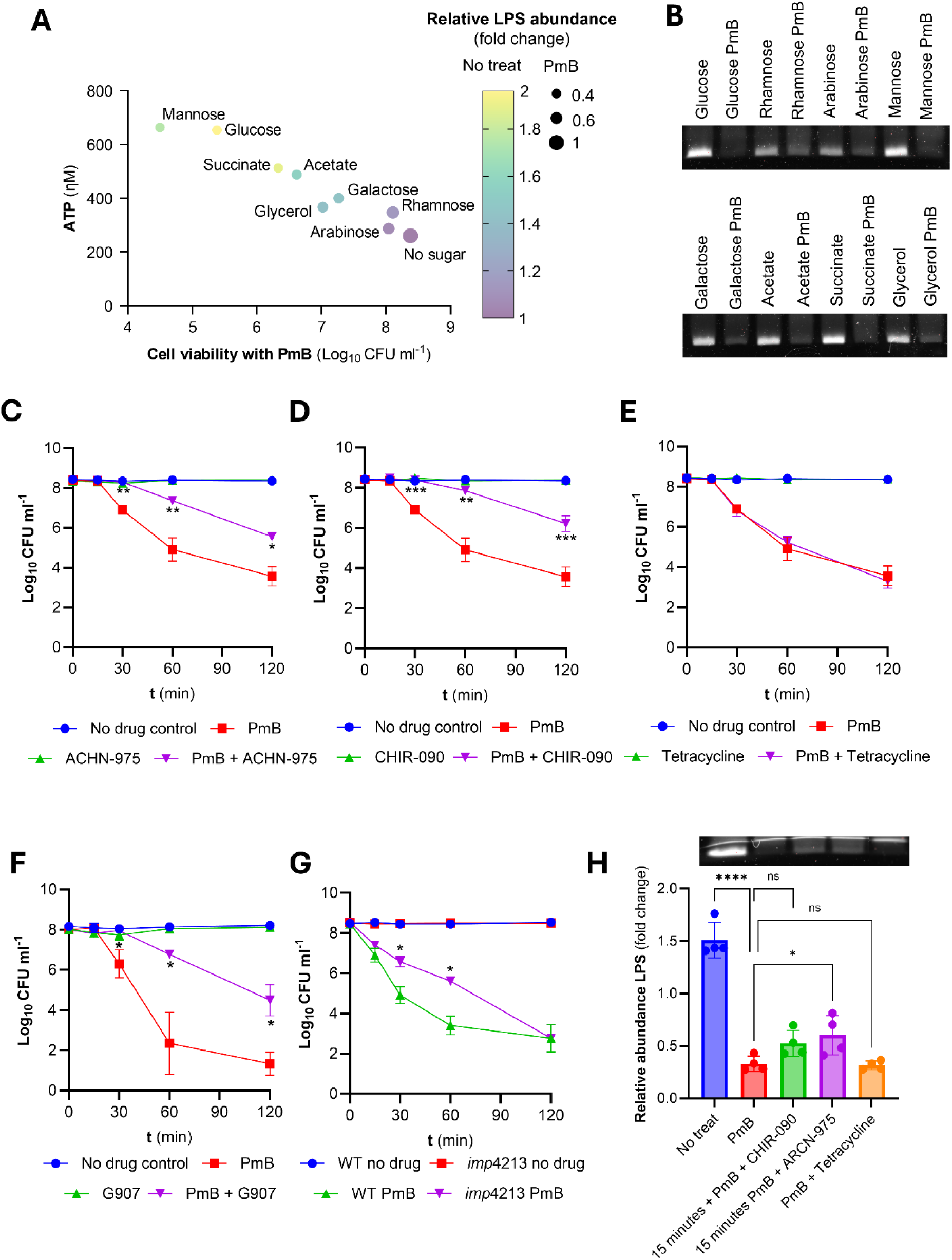
PmB-triggered LPS loss and bacterial killing require ATP and LPS synthesis and transport. **(A)** 4-way representation of the effect of various carbon sources, or none, on ATP production and LPS levels in stationary phase *E. coli* in the absence of PmB, and bacterial survival and LPS levels after exposure to 4 µg ml^-1^ PmB. The more yellow the symbol, the more LPS is produced in the absence of PmB, the smaller the symbol, the more LPS that is lost during PmB exposure. **(B)** SDS-PAGE analysis of LPS from stationary phase *E. coli* exposed to various carbon sources in MM for 15 min in the absence or presence of 4 µg ml^-1^ PmB. **(C, D, E, F)** Survival of stationary phase *E. coli* in MM+G supplemented, or not, with PmB, with or without growth-inhibitory concentrations (1 X MIC) LpxC inhibitors ACHN-975 **(C)** or CHIR-090 **(D)** or protein synthesis inhibitor tetracycline **(E)** or MsbA inhibitor G907 **(F)**. **(G)** survival of *E. coli* wild type or LptD4213 strain exposed, or not, to 4 µg ml^-1^ PmB. **(H)** LPS levels of non-treated *E. coli* or bacteria exposed to 4 µg ml^-1^ PmB +/− LpxC inhibitors (CHIR-090, ARCN-975) or tetracycline for 15 min. Panel includes an SDS-PAGE image of LPS band intensity. Of note, the experiments with G907 used *E. coli* LptD4213 to enable antibiotic ingress. All experiments were replicated in n=3 independent assays. Error bars show the standard deviation of the mean. Statistical significance of differences was determined by one-(**H**) or by two-way repeated measures ANOVA between the PmB-treated and PmB + antibiotic-treated groups (**C,D,E,F**) or between WT with PmB and *imp*4213 with PmB (G). P= *<0.05, **<0.01, ***<0.001, ****<0.0001, ns=not significant.

To establish a link between LPS production and PmB-induced LPS loss and killing, we performed PmB killing assays in the presence of various inhibitors [41]. Inhibition of LpxC, the rate-limiting step of lipid A synthesis, reduced the rate and degree of PmB-mediated killing of both *E. coli* and *P. aeruginosa*, whereas the protein synthesis inhibitor tetracycline had no effect (Fig. 3C,D,E, Supplementary Fig. S12A,B,C). There was also a slower progression of PmB-triggered surface protrusions in the presence of CHIR-090 relative to PmB alone (Supplementary Fig. S12D). We also found that an inhibitor of the LPS transporter MsbA reduced killing by PmB, whilst a strain with reduced LptD function was also killed more slowly than the wild type (Fig. 3F,G). For all inhibitors, PmB binding to the OM was not affected, as determined via NPN uptake, whereas reduction of LPS synthesis and transport significantly reduced PmB-induced IM permeation as measured by SYTOX ingress into the cytoplasm (Supplementary Fig. S13,S14).

Further evidence for the importance of LPS transport came from experiments with novobiocin, which promotes LPS trafficking from the IM to the OM [42] and was found to promote PmB-mediated LPS loss (Supplementary Fig. S15). This finding also provides a molecular explanation for the potentiating effect of novobiocin on polymyxin susceptibility [42,43].

Combined, these experiments revealed that the magnitude of PmB-mediated LPS loss correlated with metabolic activity, which in turn correlated with bacterial killing. The role of metabolic activity in LPS loss was due, at least in part, to a requirement for LPS synthesis and transport.

### Loss of LPS from the OM enables PmB-mediated disruption of the IM and bacterial killing

To differentiate between PmB action on the OM and on the IM, we examined if PmB could kill metabolically active cells by LPS loss alone, without the need for permeabilization of the IM. For this purpose, we first exposed *E. coli* to PmB for 15 min in MM+G, long enough for LPS loss to reach its maximum, but before IM disruption occurred and before CFU counts started to decrease relative to the inoculum (Fig. 1A,E, Supplementary Fig. S8). We then washed the bacteria to remove unbound antibiotic, followed by incubation in MM +/− PmB and +/− glucose. In the absence of PmB, bacteria survived at the same level over 2h in the presence or absence of glucose (Fig. 4A). By contrast, bacteria that were re-exposed to PmB were efficiently killed, regardless of the presence of glucose (Fig. 4A). Therefore, PmB-mediated LPS loss alone is not sufficient to kill bacteria. Moreover, the requirement for metabolic activity for PmB-mediated killing appeared to only apply to LPS loss from the OM and is not required for subsequent IM disruption and killing.

**Figure 4.**
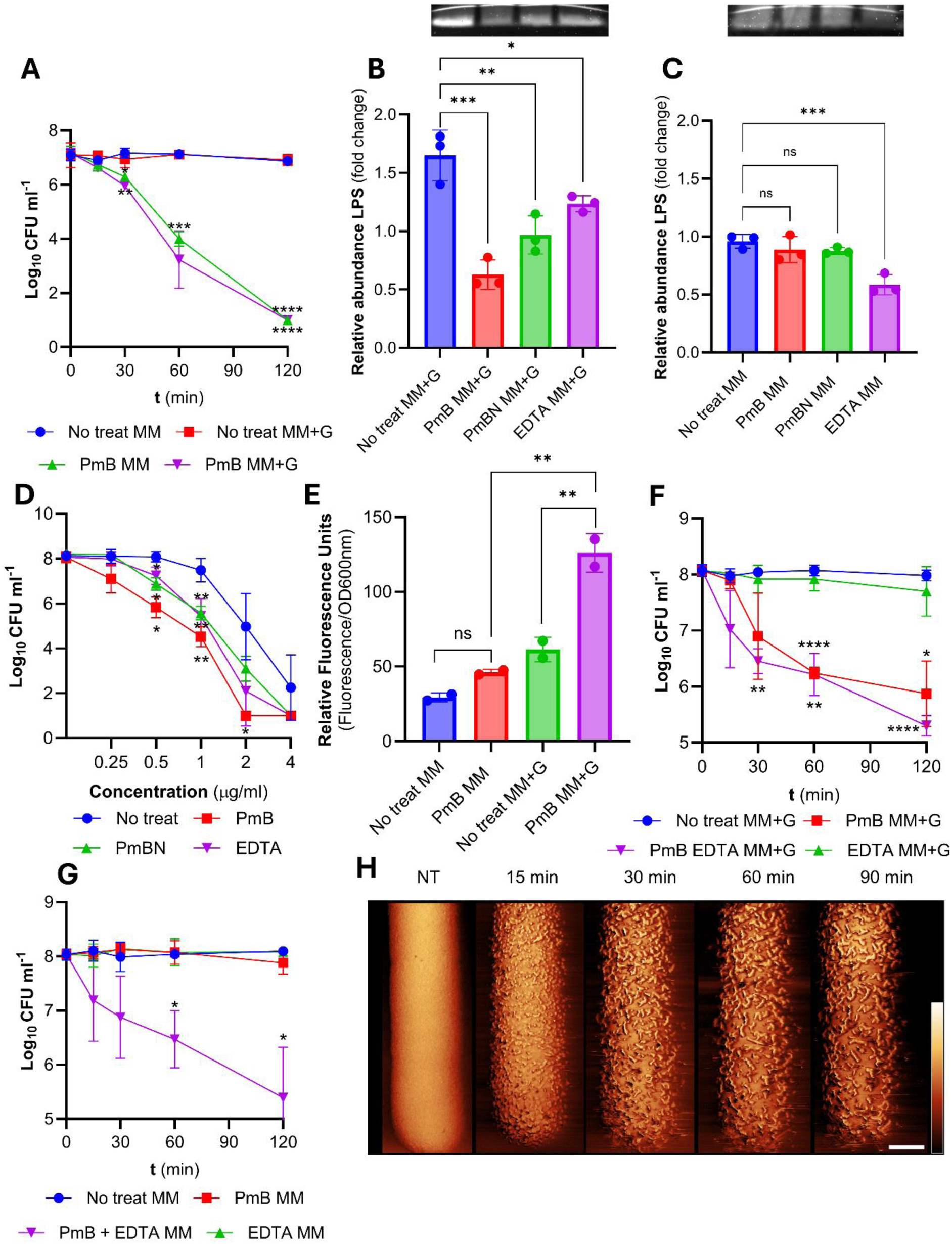
PmB-mediated LPS loss is necessary for lethality because it provides the antibiotic with access to the IM. **(A)** Survival (CFU) of stationary phase *E. coli* first pre-treated with 4 µg ml^-1^ PmB in MM+G for 15 min before washing to remove unbound antibiotic, and next (from t = 0) exposed to 4 µg ml^-1^ PmB, or not, in MM +/− glucose. **(B)** LPS levels of untreated *E. coli* or bacteria exposed to 4 µg ml^-1^ PmB, 4 µg ml^-1^ PmBN or 10 mM EDTA for 15 min in MM+G. NB: experiments with EDTA used MM that was not supplemented with MgSO_4_ or CaC^l^2. Panel includes SDS-PAGE image of LPS band intensity. **(C)** LPS levels of non-treated *E. coli* or bacteria exposed to 4 µg ml^-1^ PmB, 4 µg ml^-1^ PmBN or 10 mM EDTA for 15 min in MM. Panel includes SDS-PAGE image of LPS band intensity. **(D)** Survival of stationary phase *E. coli* exposed to a range of PmB concentrations for 2h in MM+G, following a 15-minute pre-treatment, or not, with 4 µg ml^-1^ PmB, 4 µg ml^-1^ PmBN or 10 mM EDTA in MM+G. **(E)** Levels of bodipy-vancomycin labelling of *E. coli* exposed, or not, to 4 µg ml^-1^ PmB in MM +/− glucose for 15 mins. **(F)** Survival of *E. coli* exposed, or not, to PmB with or without 10 mM EDTA in MM+G. **(G)** Survival of *E. coli* exposed, or not, to PmB with or without 10 mM EDTA in MM. **(H)** AFM phase images showing stationary phase *E. coli* exposed to 2.5 µg ml^-1^ PmB and 10 mM EDTA in MM. Scalebar: 250 nm; colour scale: 5 deg. The same cell was imaged during the experiment. All experiments were replicated in n=3 independent assays. Error bars show the standard deviation of the mean. Statistical significance of differences was determined by one-way (**B, C, E**) or two-way repeated measures ANOVA between No treat MM+G and PmB MM+G and between No treat MM and PmB MM (**A),** between No treat and pre-treatment conditions **(D)**, between No treat and antibiotic conditions **(F,G)**. P= *<0.05, **<0.01, ***<0.001, ****<0.0001, ns=not significant.

To further test the impact of LPS loss on bacterial killing, we exposed stationary phase *E. coli* to the nonlethal PmB nonapeptide (PmBN) [18], which also caused significant LPS loss within 15 min in the presence of glucose, and to EDTA, which caused LPS loss both in presence and absence of glucose (Fig. 4B,C). At the doses tested, neither EDTA nor PmBN had any effect on bacterial viability, and did not trigger membrane protrusions (Supplementary Fig. S16,S17).

Based on these findings, we hypothesised that polymyxin-triggered LPS release, while not bactericidal on its own, facilitates antibiotic entry into the periplasm, enabling PmB to access and permeate the IM and thereby kill the cell. To test this, we exposed stationary phase *E. coli* to PmB, PmBN or EDTA for 15 min in MM+G to trigger LPS loss, as above, before removing these OM disrupting agents by washing, and finally measured bacterial survival in the presence of freshly added PmB. Each of the three pre-treatments enabled killing by PmB at concentrations below those required to kill untreated control cells, indicating that LPS loss sensitises bacteria to PmB-mediated killing (Fig. 4D).

To demonstrate that PmB-triggered LPS loss causes the OM to become permeable to molecules of the size of PmB, we showed periplasmic ingress of a bodipy-conjugated analogue of the glycopeptide antibiotic vancomycin (Mw = 1723.35 g mol^-1^ vs 1301.56 g mol^-1^ for PmB) (Fig. 4E) [43]. However, minor permeabilisation of the OM, as occurs with PmB in the absence of glucose, did not permit entry of vancomycin into the periplasm (Fig. 4E).

Since EDTA caused LPS loss from stationary phase *E. coli* in the absence of glucose, (Fig. 4C), we hypothesised that EDTA would enable PmB to access the IM and thereby kill *E. coli* in the absence of energy. Firstly, in MM+G, there was a significant loss of viability of *E. coli* exposed to PmB alone and to PmB combined with EDTA, but not EDTA alone (Fig. 4F). However, levels of bacterial viability were the same for cells exposed to PmB alone and PmB-EDTA, indicating this combination was not synergistic at the concentrations used (Fig. 4F). We then repeated this experiment in MM without glucose. In keeping with our earlier findings, metabolically inactive cells were not killed by PmB alone (Fig. 4G). However, there was a significant loss of viability of metabolically inactive cells exposed to PmB and EDTA together (Fig. 4G). In addition, analysis of EDTA/PmB co-treatment of metabolically inactive cells by AFM revealed a roughening of the bacterial surface, with numerous protrusions that resembled the effects observed of PmB alone in the presence of glucose (Fig. 4H, Supplementary Fig. S18).

Together, these experiments show that metabolic activity is required for PmB lethality because it supports LPS release, which provides the antibiotic with access to the IM, the disruption of which leads to bacterial killing. However, if LPS loss occurs via a PmB-independent mechanism, such as EDTA, then there is no need for metabolic activity, demonstrating that IM disruption occurs via an energy-independent process.

### Polymyxin resistance determinant MCR-1 blocks PmB-mediated LPS release

Mobile colistin resistance (*mcr*) genes, which encode phosphoethanolamine (pEtN) transferases, are globally disseminated in the Enterobacterales [45,46]. MCR-1, which is the most common variant, selectively modifies the 4’ phosphate of lipid A targeted by PmB, reducing susceptibility to polymyxin antibiotics [47]. Previous work showed that MCR-1 protected the IM of both whole cells and spheroplasts from colistin [28,39]. However, previous studies have shown that polymyxin antibiotics can disrupt the OM of polymyxin resistant *E. coli* as evidenced by ingress of NPN or hydrophobic antibiotics such as rifampicin [14,28,29]. As such, the impact of MCR-1 on OM integrity during PmB exposure is unclear. Based on the work above, we hypothesised that pEtN-modification would contribute to polymyxin resistance by inhibiting PmB-triggered LPS release, which would block access of the antibiotic to the IM.

To test this, we used an *E. coli* transformed with an inducible *mcr*-1 construct but allowed expression to occur at basal levels (i.e. no inducer) so that we had minimal production of MCR-1 and thus only a 4-fold increase in PmB MIC above the wild type strain (Supplementary Table S1). Despite this modest change in susceptibility, MCR-1 fully protected *E. coli* from PmB at 4 µg ml^-1^ in MM+G, whereas there was a >4-log_10_ reduction in CFU counts of the wild-type (Fig. 5A). There was also a substantial reduction in the viability of *E. coli* that produced a catalytically inactive version of MCR-1 (MCR-1*) (Fig. 5A) [48]. As expected, MCR-1 reduced but did not prevent OM disruption as indicated by NPN uptake relative to the wild type strain, whereas the IM was fully protected from PmB (Fig 5B,C) [27,28,39,48].

**Figure 5.**
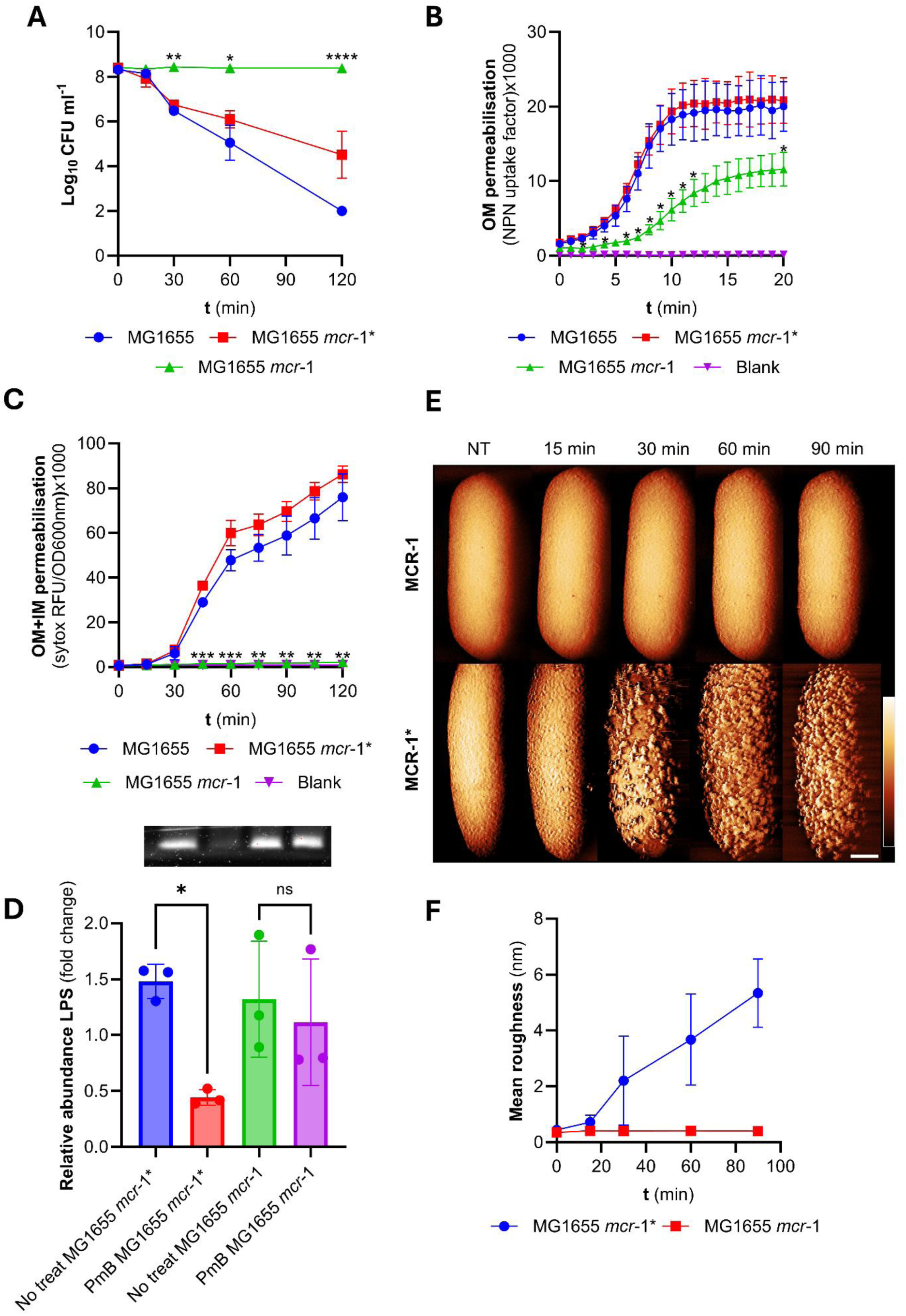
MCR-1 prevents PmB-triggered LPS release. **(A)** Survival of stationary phase MG1655, MG1655 *mcr*-1*\** or MG1655 *mcr*-1 exposed to 4 µg ml^-1^ PmB in MM+G, as determined by CFU counts. **(B)** OM disruption of stationary phase MG1655, MG1655 *mcr-*1* or MG1655 *mcr-*1 cells during the first 20 min of exposure to 4 µg ml^-1^ PmB in MM+G, as determined by uptake of NPN fluorescent dye. **(C)** OM and IM disruption of stationary phase MG1655, MG1655 *mcr-*1* or MG1655 *mcr-*1 exposed to 4 µg ml^-1^ PmB in MM+G, as determined by uptake of the fluorescent dye SYTOX green. **(D)** Total LPS in stationary phase MG1655 *mcr-*1* or MG1655 *mcr-*1 exposed, or not, to 4 µg ml^-1^ PmB in MM+G for 15 mins. Graph shows quantification of LPS levels from 3 independent experiments. **(E)** AFM phase images showing stationary phase MG1655 *mcr-*1* and MG1655 *mcr*-1 exposed to 2.5 µg ml^-1^ PmB in MM+G. The same cell was imaged during the experiment. Scalebar: 250 nm; colour scale: 4 deg. **(F)** Mean surface roughness of MG1655 *mcr*-1* and MG1655 *mcr*-1 treated with 2.5 µg ml^-1^ PmB in MM+G. All experiments were replicated in n=3 independent assays. Error bars show the standard deviation of the mean. Statistical significance of differences was determined by one-way **(D)** or two-way repeated measures ANOVA between MG1655 and *mcr-*1* or *mcr-*1 (**A, B, C**). P= *<0.05, **<0.01, ***<0.001, ****<0.0001, ns=not significant.

In support of our hypothesis, we next showed that MCR-1 prevented PmB-triggered LPS loss, as well as surface protrusions, both of which occurred in the wild type and strain expressing the catalytically inactive MCR-1 variant (Fig 5D,E,F).

Therefore, MCR-1 mediated pEtN modification of lipid A prevents LPS loss from *E. coli* exposed to PmB, which contributes to resistance by blocking access of the antibiotic to the IM. However, it is not clear if the lack of LPS release is due to reduced PmB binding, or greater membrane integrity due to the pEtN modification or a combination of both factors.

## Discussion

The mechanism(s) by which polymyxin antibiotics kill bacteria has been the focus of multiple studies, with various conclusions reached and models proposed [14,15,18,21,22,49]. In this work, we established that metabolic activity is required for bactericidal activity of a clinically relevant concentration of PmB and used this as a starting point to address some key questions in the field, including the nature and mechanism of OM disruption, how PmB accesses the IM and what happens subsequently.

Combined, our work provides a refined model for the mechanism by which polymyxin antibiotics kill bacteria (Fig. 6). We found that the initial interaction of PmB with the OM, as determined by NPN ingress, was unaffected by the metabolic state of the bacteria, which explains how polymyxins can sensitize metabolically dormant persister bacteria to antibiotics [32,51,52]. However, severe OM disruption, as indicated by loss of periplasmic mCherry, LPS loss and membrane protrusions was associated with metabolic activity, which we subsequently discovered included LPS synthesis and transport. Disruption to the OM enables PmB entry into the periplasm and access to LPS in the IM, the permeabilisation of which leads to bacterial killing.

**Figure 6.**
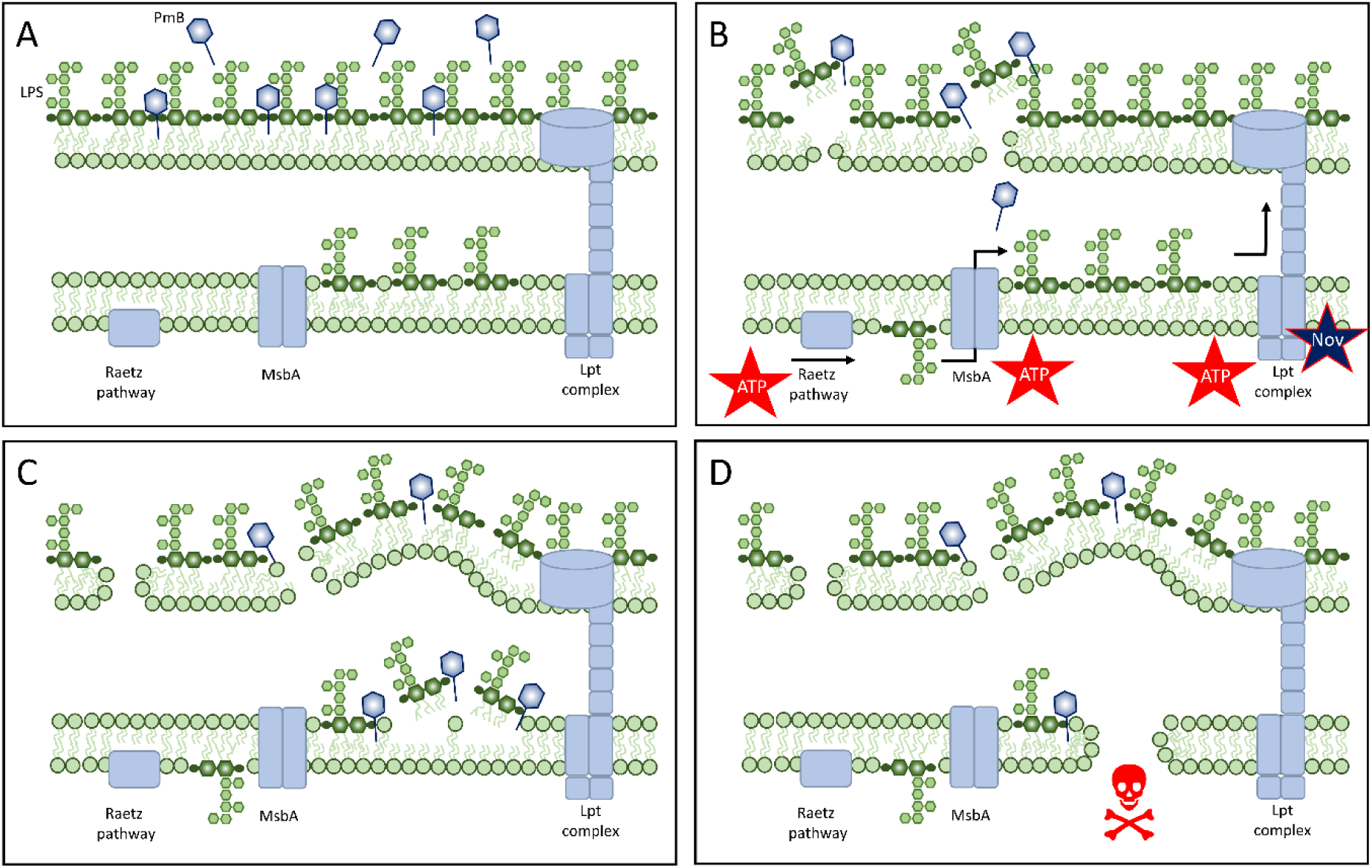
Proposed model for how PmB kills bacteria. **(A)** PmB binds to the OM, causing minor permeabilisation, regardless of bacterial metabolic activity (insufficient damage to provide PmB with access to the periplasm). **(B)** In metabolically active bacteria, PmB triggers LPS release via a process that requires ATP-dependent synthesis and transport of LPS (further boosted by novobiocin (Nov)). **(C)** The shedding of LPS compromises the integrity of the OM, resulting in surface protrusions and enables ingress of the antibiotic into the periplasm where it targets LPS in the IM, leading to permeabilisation. **(D)** The loss of integrity of both the OM and IM results in bacterial killing.

Several different classes of bactericidal antibiotics require bacterial metabolic activity for lethality [2,3,4,5]. However, our finding that this requirement also applies to PmB goes against the common assumption that the efficacy of membrane-targeting antibacterials is independent of metabolic activity. It also provides an explanation for polymyxin tolerance observed in stationary phase cells and persisters, which typically have reduced metabolic activity relative to the bulk population [4,5,9,10,31,32,53,54]. An important distinction, though, is that whilst many bactericidal antibiotics kill via metabolism-dependent production of reactive metabolites [5,8,55], the key energy-dependent step required for polymyxin activity is disruption of the OM, which enables the antibiotic to permeabilise the IM and kill the bacterium.

We have previously shown that sub-inhibitory concentrations of the LptD inhibitor murepavadin promoted polymyxin susceptibility of *P. aeruginosa*, due to accumulation of LPS in the IM without reducing LPS transport to the OM [27]. Based on the work presented here, we would predict that growth inhibitory concentrations of inhibitors of the Lpt system would reduce killing by polymyxins. However, we were unable to test this conclusively because the available inhibitors (murepavadin and thanatin) are cationic peptides that cause OM disruption, including membrane blebbing, which would make interpretation of experiments impossible [56,57,58]. Instead, we capitalised on the observation that novobiocin promoted LPS transport by enhancing ATPase activity, which in turn increased bacterial susceptibility to polymyxin antibiotics, although it was not known why at the time [42,43]. In addition to providing further evidence that LPS transport to the OM enables PmB-triggered LPS release, our work also explains how novobiocin promotes PmB activity [43].

Although our conclusions agree with some previous reports, they may appear inconsistent with others. For example, there is evidence that PmB is active against metabolically inactive bacteria. However, those studies employed either exponential phase cells that may retain a degree of metabolic activity or ATP or used assay conditions with low concentrations of nutrients or supra-physiological concentrations of polymyxins where specificity for LPS is lost due to the amphipathic nature of these antibiotics [13,15,32,33,34,35,59]. There have also been conflicting studies on the contribution of ROS to bactericidal activity of PmB, which may reflect differing experimental systems [60,61,62,63]. Our finding that energy is required for severe OM disruption does not rule out a role for ROS, which could arise via LPS synthesis, for example. Finally, whilst we show surface protrusions, it is unclear whether these are released as vesicles, which has been reported to occur at sub-inhibitory PmB concentrations [20,64,65,66,67,68]. More work is needed, therefore, to fully elucidate the origin and nature of PmB-induced vesicles and their impact on bacterial survival.

Despite the loss of LPS that occurred during PmB exposure, cells remained viable until IM disruption occurred, in agreement with our and others’ previous work showing that IM damage is required for lethality [25,27,51]. Although the OM is an important load-bearing structure, much of the mechanical strength is provided by the porin network, which remained largely intact during PmB exposure and may therefore explain how *E. coli* can tolerate major LPS loss without a reduction in viability [38,69].

The global dissemination of *mcr* genes on broad host range plasmids poses a significant threat to the clinical use of polymyxin antibiotics [45]. Intrinsic and some acquired polymyxin resistance is associated with very high levels of lipid A modification with cationic pEtN and/or L-ara4N moieties, which prevents PmB binding to the OM [14,18,20]. However, MCR-mediated resistance often involves modification of only some of the lipid A in the OM, which likely explains how PmB can bind and sufficiently permeabilise the OM for the ingress of hydrophobic antibiotics such as rifampicin [28,39]. We found that some OM permeabilisation occurred for *E. coli mcr-*1, but also showed that the MCR-1 mediated lipid A modification prevented PmB-triggered LPS release [27,28], which in turn prevented access of the antibiotic to the IM. This, coupled with the protection of the IM via LPS modification [27,28], provides distinct layers of protection against PmB.

In summary, this work advances our understanding of the mechanism of action of polymyxin antibiotics by showing that the nature and magnitude of PmB-mediated OM damage is dependent upon the metabolic activity of the cell, which in turns modulates access of the antibiotic to the IM, disruption of which is required for lethality but is not energy dependent (Fig. 6). This work also provides an explanation for how polymyxin tolerance occurs in persister and stationary phase cells, despite the membrane-disrupting activity of these antibiotics (Fig. 6).

## Methods

### Bacterial strains and growth conditions

The bacterial strains used in this study are listed in supplementary table S1. *E. coli* strains expressing *mcr*-1 or *mcr*-1* under the control of the pBAD promoter were based on those described previously and were transformed with plasmids synthesised by Thermo Fisher [48]. Unless stated otherwise, all strains were grown in Mueller-Hinton broth (MHB; Millipore) for 18 h at 37 °C with shaking (180 r.p.m.) to stationary phase. Where indicated, exponential phase bacteria were generated by inoculating fresh MHB with 1:1000 dilution of stationary phase cells followed by a second growth period of 3 h at 37 °C with shaking (180 r.p.m.) For all experimentation, bacteria were washed three times with 1X M9 minimal medium (Gibco) supplemented with 2 mM MgSO4 and 0.1 mM CaCl2 [37] (referred to here as minimal medium, MM). Washed cells were then resuspended in MM +/− 0.36% glucose (MM+G) to 10^8^ CFU ml^-1^. To enumerate bacterial CFU, 10-fold serial dilutions were made in PBS and plated onto Mueller-Hinton agar (MHA; MHB supplemented with 1.5% bacteriological agar). Inoculated agar plates were incubated statically for 18 h in air at 37°C.

### Determination of antibiotic MICs

The MIC of polymyxin B (PmB), CHIR-090, ACHN-975, tetracycline, G907, novobiocin and murepavadin against indicated bacterial strains was determined according to the well-established broth microdilution method [70]. In brief, a 96-well microtitre plate was used to prepare a range of antibiotic concentrations in 100 µl of MM+G or cation-adjusted MHB (CA-MHB) via a series of two-fold serial dilutions. For antibiotic synergy determination, checkerboard analyses were performed by preparing two-fold serial dilutions of two antibiotics, with each one across a different axis, generating an 8 x 8 matrix to assess the MICs of each antibiotic when used in combination [71]. Stationary phase bacteria were diluted to 1 x 10^6^ CFU ml^-1^ in MM+G or CA-MHB and 100 µl used to seed each well of the microtitre plate to give a final inoculation density of 5 × 10^5^ CFU ml^−1^. Plates were then incubated statically at 37°C for 18 h in air, at which point the MIC was defined as the lowest antibiotic concentration at which there was no visible growth of bacteria. In some cases, the extent of bacterial growth after incubation was also determined by obtaining OD595nm measurements using a Bio-Rad iMark microplate absorbance reader (Bio-Rad Laboratories).

### Antibiotic time-kill assays

Following three washes in MM, stationary phase or exponential phase bacteria were added at a final concentration of 1 × 10^8^ CFU ml^−1^ to 3 ml of MM, MM+G, or MM without cations and supplemented with 10 mM EDTA, containing PmB (4 µg ml^−1^) when required. This assay was also repeated in the presence of PmB and 1X MIC of a range of different antibiotics (CHIR-090, ACHN-975, tetracycline, G907), as well as in MM supplemented with a selection of equimolar concentrations of either glucose, rhamnose, arabinose, mannose, galactose, acetate, succinate, or glycerol. Cultures were incubated at 37 °C with shaking (180 r.p.m.). At each time-point (0, 15, 30, 60, and 120 min), aliquots were taken, serially diluted 10-fold in PBS and plated to determine bacterial viability by determination of CFU counts.

### Atomic force microscopy sample preparation

Bacterial imaging by AFM followed protocols described previously [37,38]. Briefly, for sample preparation, round glass coverslips of 13 mm diameter were used. They were washed by sonication in 1% w/v SDS for 10 minutes at 37kHz and 100% power, then rinsed with MQ water and ethanol, and dried with a nitrogen gun. Then they were plasma cleaned for 2 minutes at 70% power. These two procedures were repeated once more. Then coverslips were coated with Vectabond: they were placed in a beaker with a a 1:50 ratio of Vectabond solution to acetone 40 mL of acetone and 800 µL Vectabond solution for 5 minutes, then rinsed with MQ water and dried with a nitrogen gun [72]. The coverslips were glued to microscope slides with water-soluble glue purchased from Bruker (Reprorubber thin pour, Flexbar, NY). Subsequently, *E. coli* was cultured to stationary phase (18 h) in MHB, then diluted 1:100 into fresh MHB and incubated to mid-exponential phase. Bacteria were then prepared by centrifuginged 3 times at 14000 rpm for 90 seconds and resuspendinged in MM or MM+G at an OD_600_ of 0.5. (MM: 1 × M9 salts (ThermoFisher Scientific), 2 mM MgSO_4_, 0.1 mM CaCl_2_, MM+G: MM with 0.36% glucose). They were resuspended the 4_th_ time in 100 µ L 20 mM HEPES and incubated onto the Vectabond-coated glass coverslips for 5 minutes. They were then washed off with imaging media (MM or MM+G) to remove unadhered bacteria. SYTOX Green nucleic acid stain (at a final concentration of 5 µM) was added and incubated at room temperature for 5 min. We first imaged untreated bacteria in MM or MM+G, then with a pipette, this was removed as much as possible, and replaced with either PmB, EDTA, PMBN, CHIR-090 or combinations, added in the same way. Scans were subsequently taken after the introduction of the drug.

### Atomic force microscopy imaging and analysis

All experiments were performed in dynamic (AC) mode with a Nanowizard III AFM with UltraSpeed head (Bruker AXS), with an Andor Zyla 5.5 USB3 fluorescence camera on an Olympus IX 73 inverted optical microscope. FastScanD cantilevers were used, with a resonant frequency around 110 kHz and a spring constant of 0.25 N/m. The drive frequency used ranged from 95 to 120 kHz, depending on the cantilever resonance measured by calibration. The setpoint was between 10 and 15 nm (approx. 50 to 70% relative to free amplitude). The whole-cell images were acquired at 2 to 2.5 Hz line frequencies, 2 x 2 µm^2^, and 512 x 512 or 512 x 256 pixels. Higher-magnification images were acquired over 500x 500 nm^2^ and 512 x 51221 pixels, recorded at 4 to 5 Hz line frequencies. Images in figures 2A were acquired over 200x 200 nm^2^ and 512 x 512 pixels recorded at 8 to 10 Hz lie frequency..The OMV images in Fig. 2B are 800 x 800 nm^2^ and 512 x 512 pixels.  To process the images obtained from the AFM the software Gwyddion 2.65 was used (https://gwyddion.net/) [73].

The following steps were taken to obtain a post-processed image: (1) level data by mean plane subtraction; (2) level by flattening base; (3) if there were strongly protruding features, a mask was applied by selecting features above a 50% threshold, to prevent these from biasing the background correction and row alignment; (3) align rows by 2^nd^ polynomial line-by-line background subtraction (and exclude the masked region if applicable); (4) apply a 1-2px Gaussian filter to the whole image to remove high frequency noise (remove mask if applicable); (5) define zero reference and adjust the colour scale to highlight key features.

### Scanning electron microscopy

The *E. coli* cells were cultured in MHB to stationary phase before bacteria were collected using centrifugation, and the cell pellet was washed three times with M9 medium without glucose. The cells were then diluted to 10^8^ CFU ml^-1^ in M9 +/− glucose with 4 µg ml^-1^ PmB and samples were collected at various time points through centrifugation for further analysis. The samples obtained were treated with 2.5% glutaraldehyde in a 0.1 M sodium cacodylate buffer (pH=7.3) and were allowed to fix overnight at 4°C. After fixation, the cells were suspended in the 0.1 M sodium cacodylate buffer and subjected to dehydration using ethanol concentrations of 10%, 20%, 30%, 40%, 50%, 70%, 80%, and 100%, with a 10 min incubation at each step. The drying procedure was performed via Critical Point Drying, and a platinum coating of 10 nm thickness was applied after the samples were completely dried. The prepared samples were examined under a scanning electron microscope (Zeiss Crossbeam 550 FIB-SEM) at an accelerating voltage of 2 kV, and all images were analysed after conversion for differences in magnification. The ImageJ/Fiji software was used in SEM image processing and analysis.

### OM permeability assay by NPN uptake

To detect OM damage by PmB, the well-established NPN uptake assay was performed [39]. The fluorescent probe *N*-phenyl-1-naphthylamine (NPN; Acros Organics), at a final concentration of 10 μM, was added to 100 µl of 2 × 10^8^ CFU ml^−1^ stationary phase or exponential phase bacteria in MM, MM+G, or MM without cations and supplemented with 10 mM EDTA in a black microtitre plate with clear-bottomed wells (Greiner Bio-One). Relevant concentrations of PmB or PmB in combination with 1X MIC of various antibiotics were added to the wells of the microtitre plate to give a total volume of 200 µl. Fluorescence, at 37 °C with shaking, was measured immediately using a Tecan Infinite 200 pro plate reader using an excitation wavelength of 355 nm and an emission wavelength of 405 nm. Fluorescence measurements were obtained every min for 20 min, and the degree of OM permeabilisation, referred to as the NPN Uptake Factor, was calculated using the following formula:

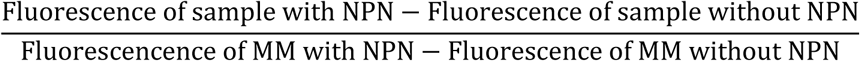

The resulting NPN Uptake Factor values were normalised according to OD_600nm_.

### OM permeability assay by mCherry egress

To further assess OM damage by PmB, _Peri_mCherry *E. coli* was prepared by transforming the pPerimCh plasmid [40] into *E. coli* MG1655. This plasmid encodes a constitutively expressed mCherry construct that accumulates in the periplasm. Stationary phase bacteria (2 × 10^8^ CFU ml^−1^) were incubated with PmB in MM or MM+G. At each timepoint, 200 µl of the culture was centrifuged to separate bacteria and culture medium, the supernatant was removed and placed in the well of a black clear-bottomed 96-well plates (Greiner Bio). The fluorescence was measured with an excitation wavelength of 587nm and an emission wavelength of 610 nm.

### IM permeability assay

To measure IM permeabilisation by PmB, the well-established SYTOX green assay was used [74]. Following washes in MM, stationary phase bacteria at an inoculum density of 1 × 10^8^ CFU ml^−1^ was added to 3 ml of MM or MM+G containing the relevant antibiotics. The fluorescent probe SYTOX green (Invitrogen) was added to these cultures at a final concentration of 1 µM. Aliquots (200 μl) were transferred to a black microtitre plate with clear-bottomed wells (Greiner Bio-One). Fluorescence and OD_600nm_ were measured in a Tecan Infinite 200 Pro plate reader (excitation at 535 nm, emission at 617 nm) every 15 min for 2 hr at 37 °C with shaking. Raw fluorescence readings were normalised according to OD_600nm_.

### Determination of Lipid A modification by mass spectrometry

Lipid A was prepared and analysed for modifications by mass spectroscopy as described previously [27,28]. Briefly, Lipid A was detached from carbohydrates via mild acid hydrolysis (2% acetic acid, 100 °C 30 min), washed in H_2_O and loaded on the target and overlaid with 9H-Pyridol[3,4-B]indole (Norharmane, Sigma-Aldrich) used at 10 mg ml^-1^ in Chloroform:Methanol 9:1 and mixed before air drying and MALDI-TOF analysis using a MALDI Biotyper Sirius system (Bruker Daltonics, USA).

### Detection of LPS by SDS-PAGE

The experimental conditions for LPS quantification were the same as those used for the antibiotic time-kill assay. Briefly, 1 × 10^8^ CFU ml^−1^ of stationary phase bacteria were added to 3 ml cultures of MM or MM+G in the presence of either PmB, PmB with 1X MIC of various antibiotics, PmBN, or EDTA. LPS quantification was also performed in the presence of PmB in MM supplemented with a selection of equimolar concentrations of either glucose, rhamnose, arabinose, mannose, galactose, acetate, succinate, or glycerol. Following 15 min incubation at 37 °C with shaking (180 r.p.m.), samples were pelleted at 16000 *g* and washed twice with PBS. Pellets were incubated with 2% SDS and DNAse1 and incubated for 10 min at room temperature. the pellet was resuspended in 100 µl of 1X sample buffer (4% β-mercaptoethanol, 10% Glycerol, 0.1 M Tris, 2% SDS of a pH 6.8). The volume of sample buffer was adjusted according to the fold-change in OD_600nm_ compared to time point zero to give equal biomass between samples. Next, samples were incubated for 10 min at 95°C. They were then checked to contain equal biomass by separation on a 12% Tris-glycine gel, followed by Coomassie staining. The samples were then incubated with 100 µg ml^−1^ proteinase K overnight at 55°C to remove protein and again separated on a 12% Tris-glycine gel. The gel was fixed overnight in a solution of 50% methanol, and 5% acetic acid. LPS was then quantified using the Pro-Q™ Emerald 300 Lipopolysaccharide Gel Stain Kit (Invitrogen) according to the manufacturer’s protocol. Densitometric analysis was performed using Fiji ImageJ with the resulting values used to interpolate the standard curve performed in Supplementary Fig 8C, to calculate relative LPS abundance.

### Detection and quantification of released LPS

This protocol is provided by Dr Robert Hancock (https://cmdr.ubc.ca/bobh/method/kdo-assay/). Briefly, 50 µL of supernatant (generated from a 15 min incubation of 1 × 10^8^ CFU ml^−1^ of stationary phase bacteria in MM +/−glucose and +/− 4 µg mL^-1^) was boiled in 50 µL 0.5N H_2_SO_4_ for 15 min, followed by the sequential addition of 200 µL arsenite reagent (0.3 µM NaAsO_2_ dissolved in 0.5N HCL), 800 µL 42 mM 2-thiobarbituric acid. The solution was boiled for a further 10 min before the addition of 1.5 mL butanol reagent (5.0 mL of concentrated HCl added to 95 mL n-butanol). The butanol layer was separated from the solution by centrifugation for 5 min at 2000 rpm. The Kdo extinction coefficient (OD_552nm_ – OD_509nm_) was calculated by measuring 200 µL for absorbance at OD552nm and OD509nm in a Tecan Infinite 200 Pro plate reader. The difference in OD was used to interpolate a standard curve generated by performing Kdo analysis on a 1/10 serial dilution of 5 mg mL^-1^ of purified rough LPS (Supplementary Fig 8A).

### ATP quantification

To quantify ATP levels of stationary phase bacteria in different sugar conditions, an ATP standard curve was first performed using the BacTiter-Glo™ (Promega) kit. A 10-fold serial dilution series of ATP was performed in MM over a concentration range of 1000-0.01 nM in a white microtitre plate (Greiner Bio-One). The BacTiter-Glo™ buffer was added to the substrate to make the BacTiter-Glo™ substrate. This substrate was combined with the dilution series and luminescence was measured in a Tecan Infinite 200 Pro plate reader using a 100 ms integration time. Following this, stationary phase bacteria at an inoculum density of 1 × 10^8^ CFU ml^−1^ was added to MM or MM containing a range of equimolar sugar conditions. At each time point (0, 15, 30 min), 100 µl of the culture was combined with 100 µl of the substrate in a white microtitre plate and luminescence measured. ATP concentrations under these different conditions were interpolated from the standard curve.

### Measuring vancomycin binding

Following three washes in MM, stationary phase bacteria were added at a final concentration of 1 × 10^8^ CFU ml^−1^ to 3 ml of MM +/− glucose and +/− 4 µg mL^-1^ PmB, in the presence of 10 µg mL^-1^ BODIPY™ FL Vancomycin (Invitrogen). Cultures were incubated at 37°C for 15 min, followed by 3 washes in PBS to remove unbound BODIPY™ FL Vancomycin. Cultures were resuspended in 3 ml PBS and 200 µL added to a black clear-bottomed 96-well plates (Greiner Bio). Fluorescence and OD_600nm_ were measured in a Tecan Infinite 200 Pro plate reader (excitation at 480 nm, emission at 520 nm). Raw fluorescence readings were normalised according to OD_600nm_.

### Statistical analysis

Experiments were performed on at least three independent occasions, and the resulting data are presented as the arithmetic mean of these biological repeats unless stated otherwise. Error bars, where shown, represent the standard deviation of the mean. For single comparisons, a two-tailed Student’s *t*-test was used to analyse the data. For multiple comparisons at a single time point or concentration, data were analysed using a one-way analysis of variance (ANOVA) or a Kruskal–Wallis test. Where data were obtained at several different time points or concentrations, a two-way ANOVA was used for statistical analyses. Appropriate post hoc tests (Dunnett’s, Tukey’s, Sidak’s, Dunn’s) were carried out to correct for multiple comparisons, with details provided in the figure legends. Asterisks on graphs indicate significant differences between data, and the corresponding p-values are reported in the figure legend. All statistical analyses were performed using GraphPad Prism 7 software (GraphPad Software Inc, USA).

## Acknowledgements

Steven Rutherford and Kerry Buchholz (Genentech), Thomas Clarke and Simon Stoneham (Imperial College London) and Georgina Benn (University College London/University of Oxford) are thanked for helpful discussions.

## Author contributions (order TBD based on author order)

C.B., E.J.A.D, W.J.L., B.B.B., A.M.E. and B.W.H. designed research; C.B., E.J.A.D, S.M.A.R., A.E.L., W.J.L., G.L.M. and A.M.E. performed research; C.B., E.J.A.D, S.M.A.R., A.E.L., W.J.L., A.M.E. and B.W.H. analyzed data; C.B., E.J.A.D, B.B.B., A.M.E. and B.W.H. wrote the paper.

