## Supplementary data file for "Polymyxin B lethality requires energy-dependent outer membrane disruption"

This file contains supplementary table 1 and supplementary figures 1-18

| **Species/Strain** | **Relevant characteristics and source** | **PmB MIC ($\mu$g ml$^{-1}$)** |
|---|---|---|
| *E. coli* MG1655 | Well characterised K-12 laboratory strain [75] | 0.25 |
| | P*mcr-1* [This study] | 1* |
| | P*mcr-1** [This study] | 0.25 |
| *E. coli* MC4100 | Well characterised K-12 laboratory strain [76] | 0.25 |
| *E. coli imp*4213 | Impaired OM barrier function due to a deletion in *lptD* [77,78] | 0.125 |
| *E. coli* CFT073 | Uropathogenic clinical isolate [79] | 0.25 |
| *E. coli* KPC BM16 | Clinical isolate [28,80] | 0.5 |
| *E. coli* DIN | Clinical isolate [28,80] | 0.25 |
| *E. coli* ATCC 25922 | Type strain used as a reference for antibiotic susceptibility testing | 0.25 |
| *K. pneumoniae* IMP12 | Clinical isolate expressing IMP carbapenemase [81] | 0.5 |
| *K. pneumoniae* IMP76 | Clinical isolate expressing IMP carbapenemase [81] | 1 |
| *C. freundii* IMP61 | Clinical isolate expressing IMP carbapenemase [81] | 0.5 |
| *E. asburiae* IMP8 | Clinical isolate expressing IMP carbapenemase [81] | 1 |
| *P. aeruginosa* PA14 | Well characterised laboratory strain [82] | 0.125 |
| *P. aeruginosa* AK3 | Isolate from person with cystic fibrosis. Small colony variant phenotype [27]. | 0.5 |
| *P. aeruginosa* AK11 | Isolate from person with cystic fibrosis. Mucoid phenotype [27]. | 0.125 |
| *A. baumannii* AS | Clinical isolate [This study] | 0.25 |
| *A. baumannii* N12 | Clinical isolate [This study] | 0.25 |

*For MG1655 P*mcr*-1, the uninduced MIC value is provided.

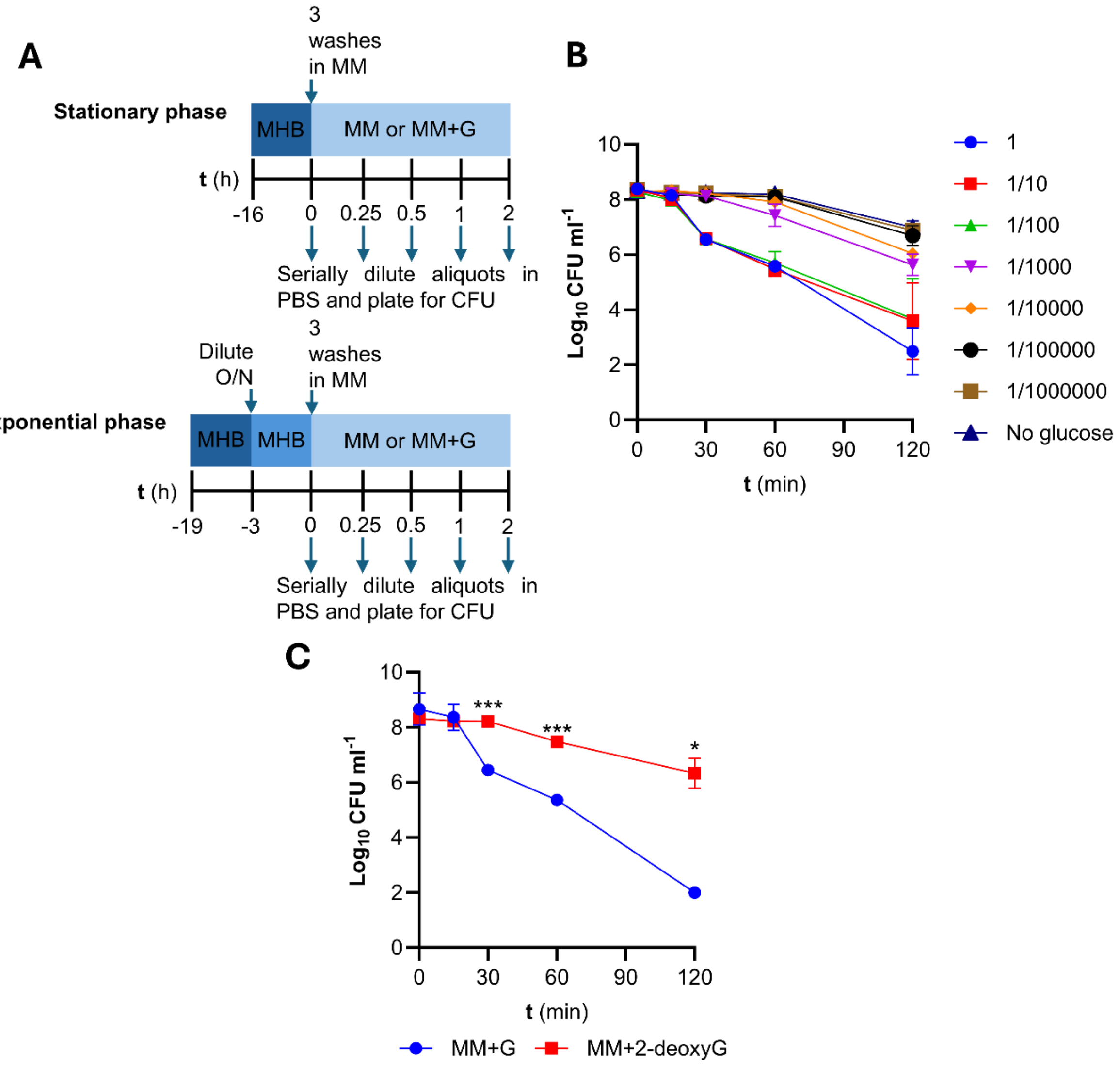


**Supplementary Figure S1. Polymyxin B lethality requires metabolic activity. (A)** Visual representation of the set-up for all microbiological assays. *E. coli* MG1655 was grown in 3 ml of MHB for 16 h at 37°C 180 r.p.m. to stationary phase. The culture was washed three times in MM and resuspended at an inoculum density of $1x10^8$ in 3 ml of MM or MM+G. To test exponential phase *E. coli,* the overnight culture was diluted 1/1000 in fresh MHB and grown for a further 3 h. **(B)** Survival of stationary phase *E. coli* exposed to 4 µg $ml^{-1}$ PmB from t = 0 in the presence of a 1/10 dilution series of 0.36% glucose. A 1/100 dilution of 0.36% glucose supported PmB killing, however, a 1/1000 dilution substantially reduced the rate and degree of PmB killing. **(C)** Survival of stationary phase *E. coli* exposed to 4 µg $ml^{-1}$ PmB from t = 0 in MM+G or MM with an equimolar concentration of 2-deoxy-D-glucose, as determined by CFU counts**.** All experiments were replicated in n=3 independent assays. Error bars show the standard deviation of the mean. Significant differences were determined by two-way repeated measures ANOVA. P= *<0.05, ***<0.001.

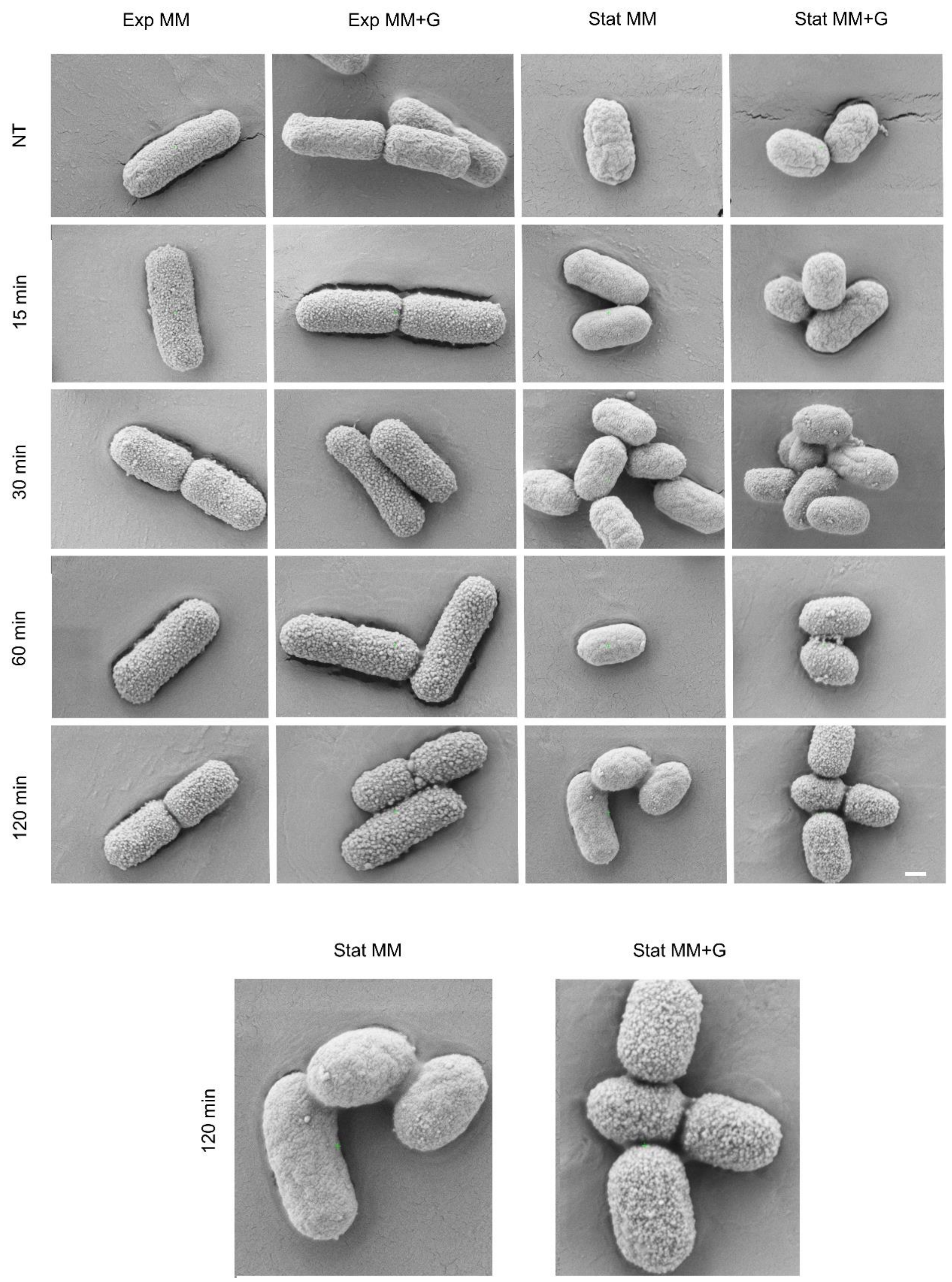


**Supplementary Figure S2. PmB causes surface protrusions on exponential phase *E. coli* MG1655 and stationary phase cells when glucose is present.** Exponential (Exp) or stationary-phase (Stat) *E. coli* cells were exposed to PmB (4 µg $ml^{-1}$) in MM +/- glucose and samples taken at the indicated time points. In keeping with the data from AFM studies, stationary phase cells exposed to PmB in the presence of glucose began to show surface protrusions from ~30 min, which increased over time. Conversely, stationary phase cells exposed to the polymyxin without glucose showed minimal to no surface protrusions. PmB triggered surface protrusions in exponential phase cells whether glucose was present or not. Images for stationary phase cells at 120 min are shown again in an enlarged image to highlight differences in surface appearance. Scale bar represents 400 nm.

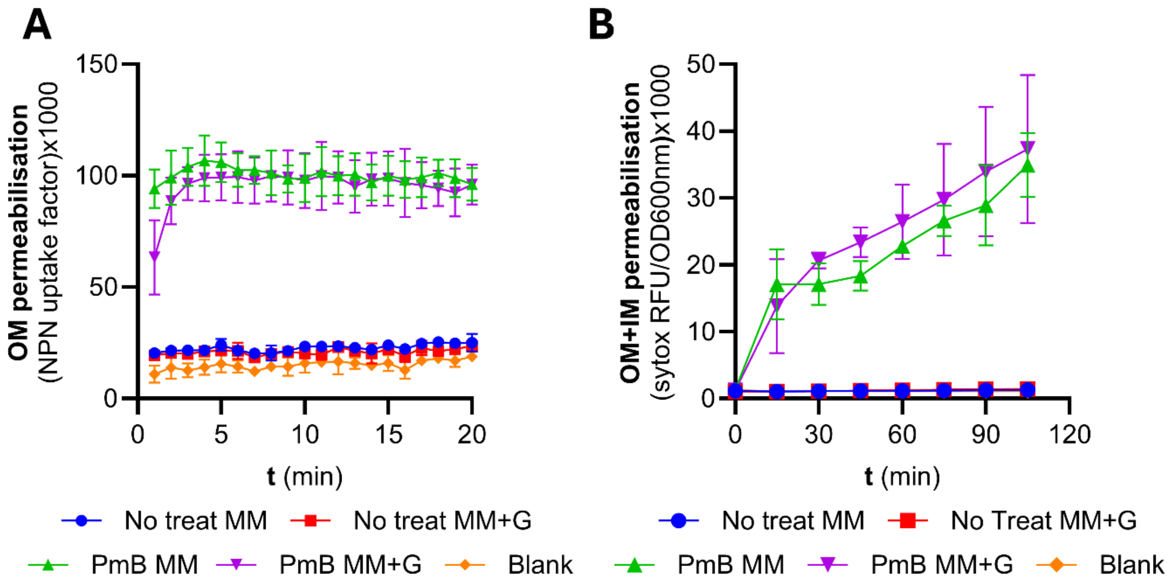


**Supplementary Figure S3. Metabolic activity does not affect the ability of PmB to disrupt the OM and IM of exponential phase *E. coli.* (A)** OM disruption of exponential phase *E. coli* cells during the first 20 min of exposure to 4 µg $ml^{-1}$ PmB in MM +/- glucose, as determined by uptake of the NPN fluorescent dye. **(B)** OM and IM disruption of exponential phase *E. coli* exposed to 4 µg $ml^{-1}$ PmB in MM +/- glucose, as determined by uptake of the fluorescent dye SYTOX green. All experiments were replicated in n=3 independent assays. Error bars show the standard deviation of the mean. No significant differences were observed between exponential phase *E. coli* in MM +/- glucose for either NPN or SYTOX uptake.

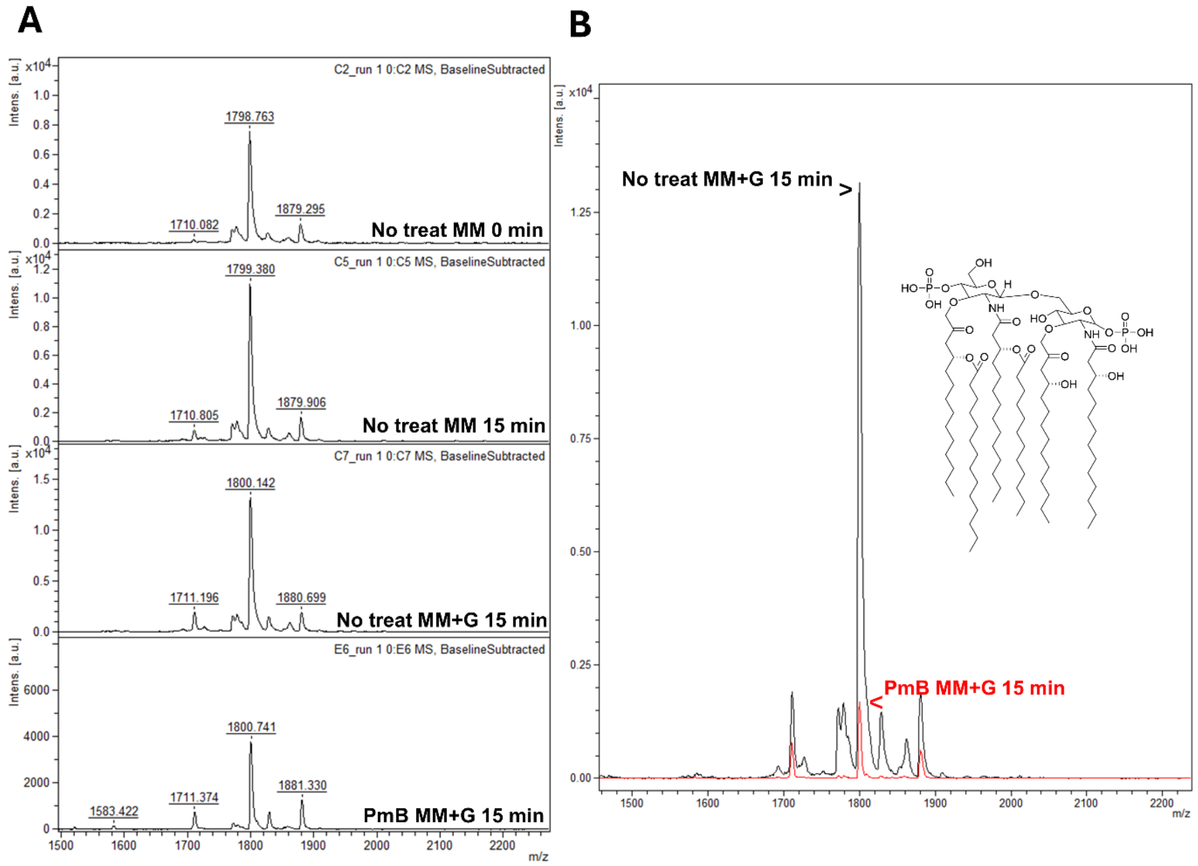


**Supplementary Figure S4. Tolerance to PmB is not associated with modifications to lipid A. (A)** From top to bottom, representative mass spectra showing unmodified lipid A from stationary phase *E. coli* at time 0 min in MM, stationary phase *E. coli* at time 15 min in MM, and stationary phase *E. coli* at time 15 min in MM+G, and stationary phase *E. coli* at time 15 min in MM+G exposed to 4 µg $mL^{-1}$ PmB. The peak at ~1800 *m/z* corresponds to native hexa-acyl diphosphoryl lipid A containing four C14:0 3-OH, one C14:0 and one C12:0 [83]. The peaks at ~1710 and ~1800 *m/z* correspond to the subtraction or addition of phosphate to the native form of lipid A. **(B)** Overlaid mass spectra of stationary phase *E. coli* at time 15 min in MM+G (black) and stationary phase *E. coli* at time 15 min in MM+G exposed to 4 µg $mL^{-1}$ PmB (red), indicating a substantial reduction in native lipid A.

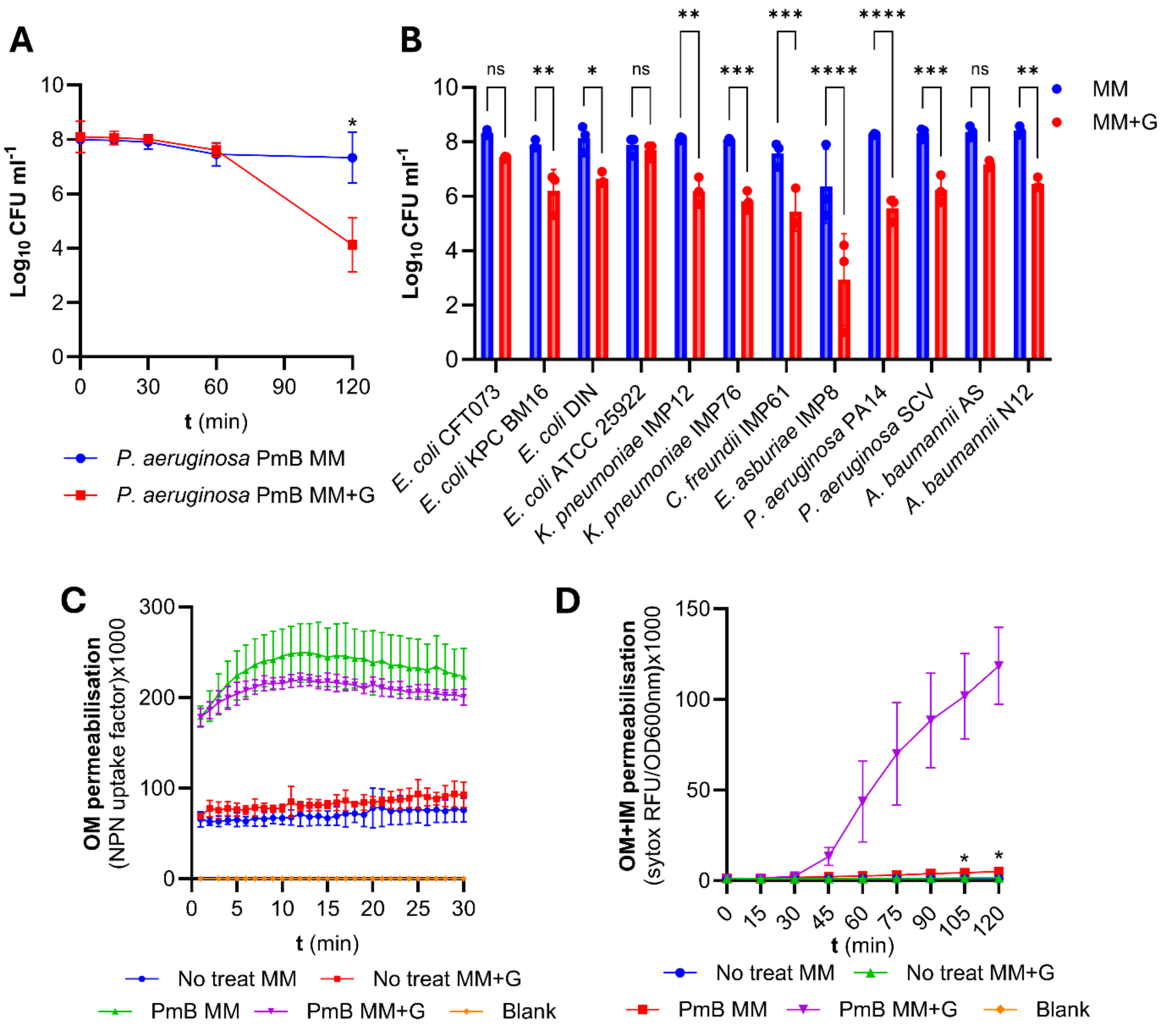


**Supplementary Figure S5. Tolerance to PmB is conserved under nutrient-limiting conditions. (A)** Survival of stationary phase *Pseudomonas aeruginosa* PA14 exposed to 4 µg $ml^{-1}$ PmB in MM +/- glucose, as determined by CFU counts. **(B)** Survival of a panel of clinical isolates cultured to stationary phase, exposed to 4 µg $ml^{-1}$ PmB in MM +/- glucose for 2 h. **(C)** OM disruption of stationary phase *P. aeruginosa* cells during the first 20 min of exposure to 4 µg $ml^{-1}$ PmB, as determined by uptake of the NPN fluorescent dye. **(D)** OM and IM disruption of stationary phase *P. aeruginosa* exposed to 4 µg $ml^{-1}$ PmB in MM +/- glucose, as determined by uptake of the fluorescent dye SYTOX green. All experiments were replicated in n=3 independent assays, error bars show the standard deviation of the mean. Significant differences were determined by one-**(B)** or two-way **(A, C, D)** repeated measures ANOVA. P= *<0.05, **<0.01, ***<0.001, ****<0.0001, ns=not significant.

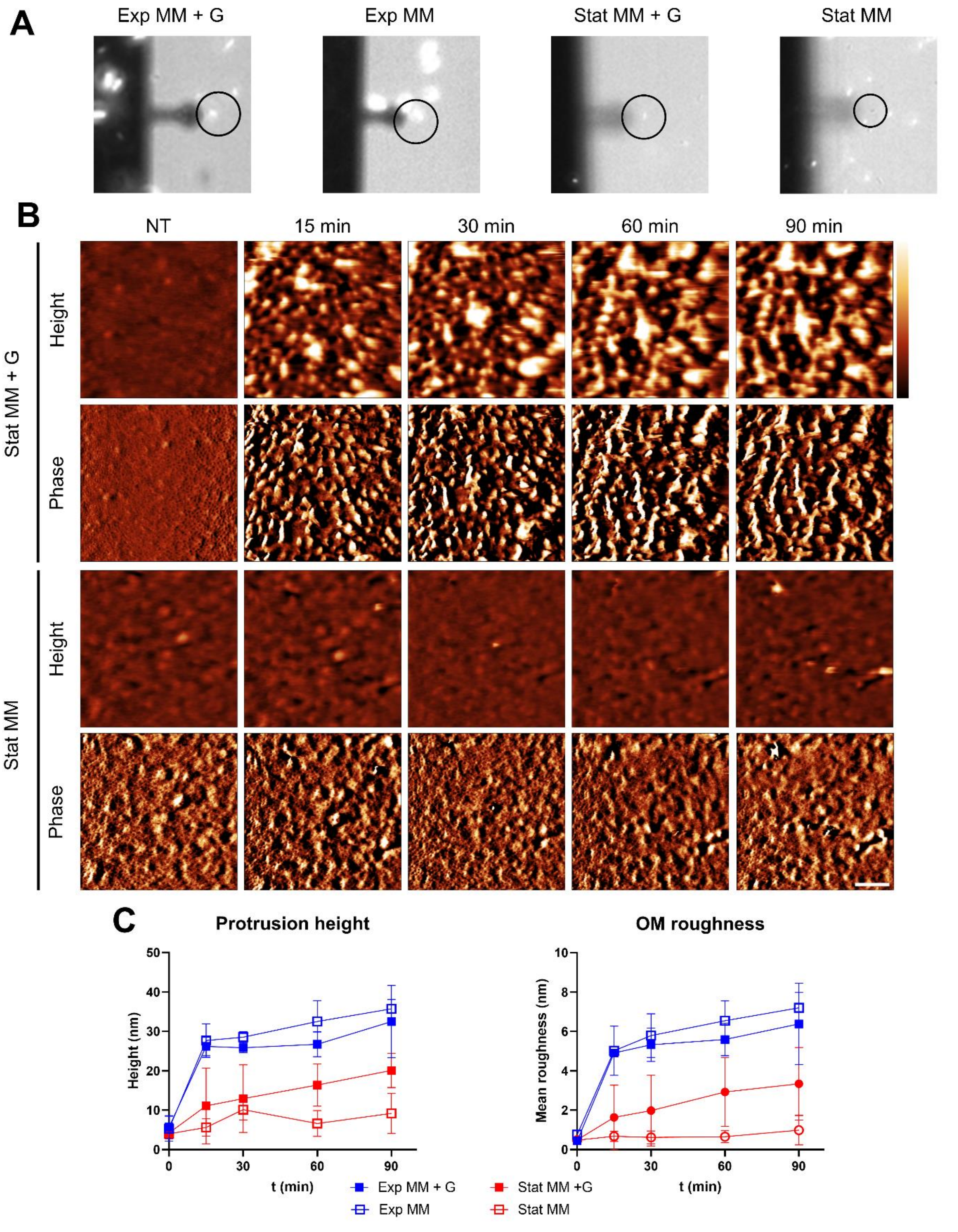


**Supplementary Figure S6. PmB leads to morphological changes to the OM at the nanoscale. (A)** Combined brightfield and fluorescence (SYTOX) images of the AFM scan region for the experiments in Fig. 1B with exponential (Exp) and stationary (Stat) phase *E. coli* at 90 min post PmB treatment (the circled cells are those chosen for the image sequences shown here and were all SYTOX-positive by 90 mins, except for stationary phase *E. coli* in MM). **(B)** Higher-magnification AFM height and phase scans of stationary phase *E. coli*. The surface of untreated cells is covered in a network of pores, here best visible in the phase images. The resolution of pores (trimeric OMPs) is ultimately compromised by the progressive roughening of the OM after PmB is added in the presence of glucose (MM + G). In MM + G, PmB caused disruption to the OM in the form of protrusions. Without glucose the appearance of the OM at the nanoscale did not change. **(C)** Quantification of protrusion height by taking the median height of features above 50% of the maximum height of the image. **(B)** OM mean roughness computed by Gwyddion. Measurements were taken from 3 different 500 nm scans of 3 different *E. coli* MG1655 cells imaged in separate experiments. Scalebar: 100 nm; colour bars: height 20 nm, phase 2 deg (row 2) and 1 deg (row 4).

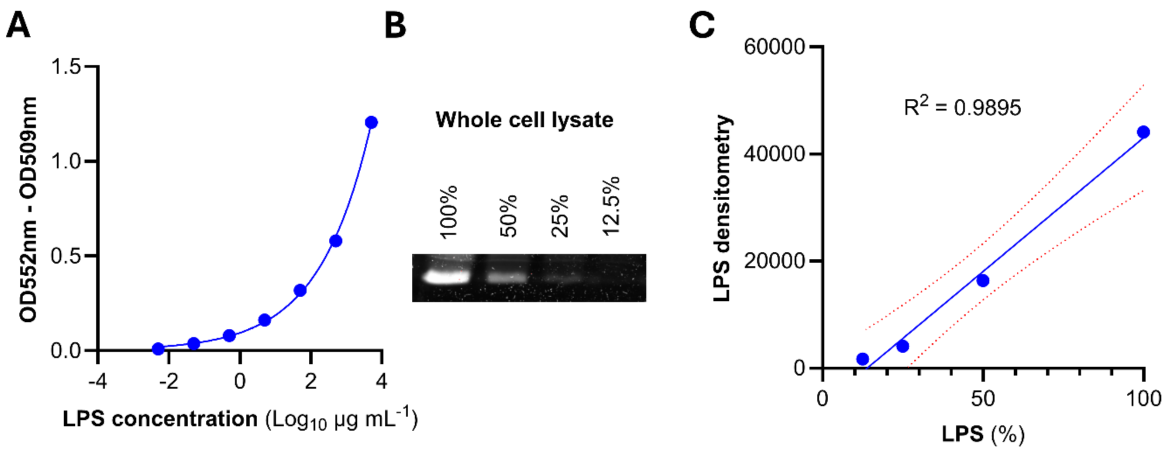


**Supplementary Figure S7. The detection limit for Kdo analysis and Pro-Q™ Emerald 300 Lipopolysaccharide Gel Stain Kit. (A)** Kdo quantification was performed by acid hydrolysis, followed by reacting with sodium arsenite and thiobarbituric acid, yielding a coloured product. A standard curve plotting a 1:10 serial dilution series of purified rough LPS against Kdo extinction coefficient (OD552nm – OD509nm) was used to interpolate released LPS in the supernatant of stationary phase *E. coli* exposed, or not, to 4 µg $ml^{-1}$ PmB in MM +/- glucose for 15 mins. Sigmoidal 4PL regression analysis was performed using prism version 10.4.1 **(B)** The detection limit of the Pro-Q™ Emerald 300 LPS gel stain kit was determined by performing a 1:2 dilution series of stationary phase *E. coli* cells incubated in MM + glucose for 15 min. **(C)** Following densitometric analysis of this dilution series, a standard curve plotting whole cell lysate LPS % against densitometry was produced. Simple linear regression was performed using prism version 10.4.1, where the blue line represents the line of best fit and the dotted red lines the 95% confidence interval.

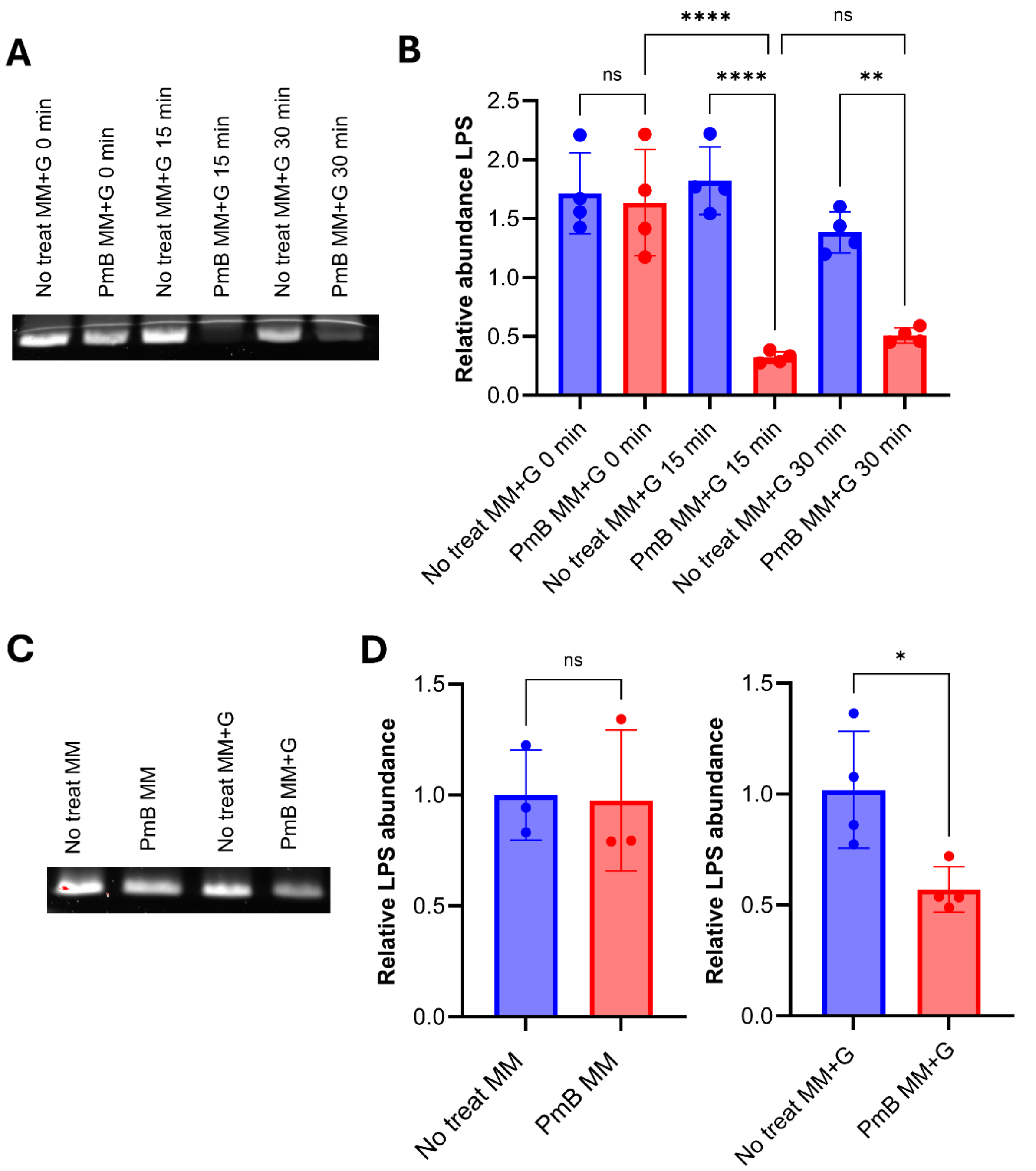


**Supplementary Figure 8. PmB-induced LPS loss is maximal at 15 minutes and is a conserved phenotype. (A)** Levels of total LPS in *E. coli* exposed, or not, to 4 µg ml$^{-1}$ PmB for 0-, 15-, and 30 min in MM+G. **(B)** Bar graph of LPS levels according to densitometry analysis of (A). Densitometric values were interpolated according to the standard curve in Supplementary Fig S7C and subsequently normalized to the No treatment controls, to give 'Relative abundance LPS'. **(C)** Levels of total LPS in *P. aeruginosa* exposed, or not, to 4 µg ml$^{-1}$ PmB in MM +/- glucose. **(D)** Bar graph of LPS levels according to densitometry analysis of (**A**). All experiments were replicated in n=3 independent assays. Error bars show the standard deviation of the mean. Significant differences were determined by one-way ANOVA P= **<0.01, ****<0.0001, ns=not significant.

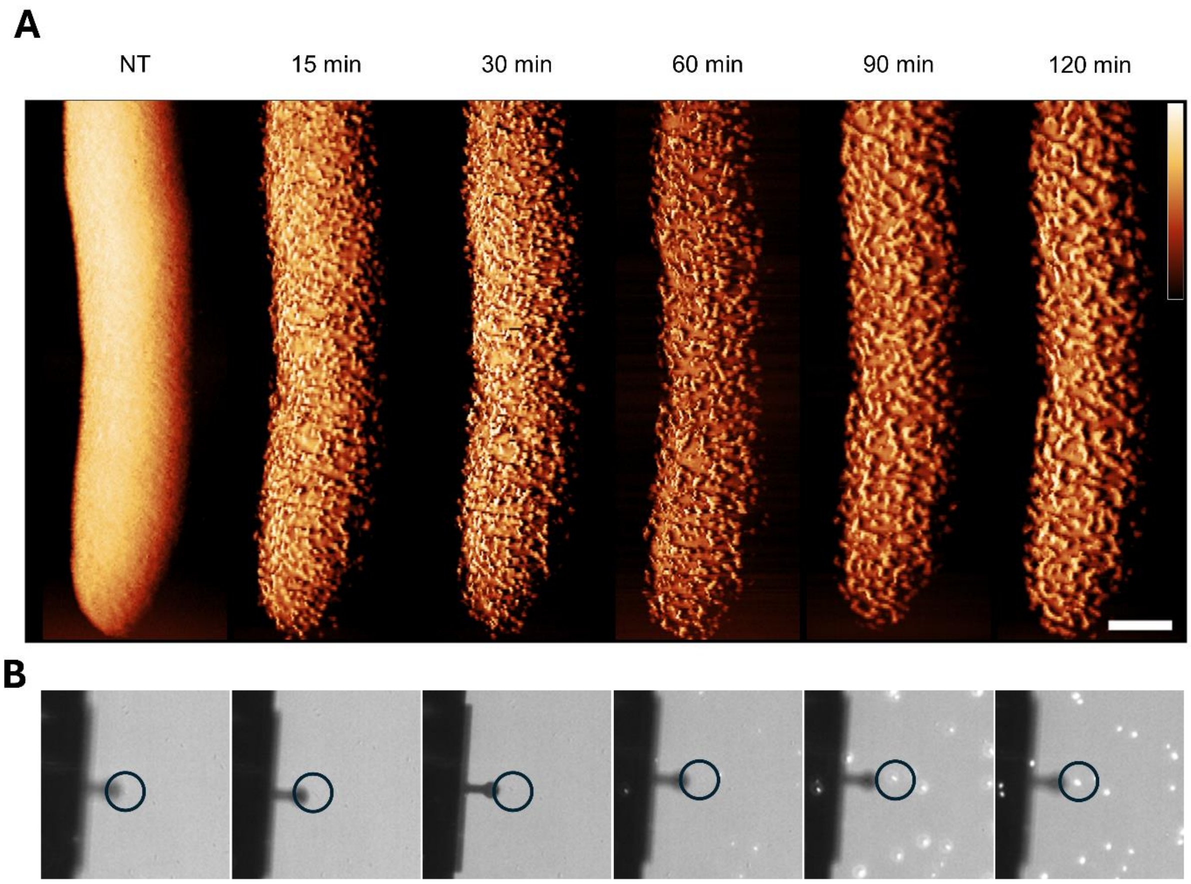


**Supplementary Figure S9. (A)** AFM phase images showing stationary phase *E. coli* cells exposed to 2.5 µg $ml^{-1}$ PmB in MM + G, shown as a function of time. Scalebar: 250 nm. Colour scale: 9 deg. **(B)** Combined brightfield and fluorescence (SYTOX) images of the AFM scan region for the experiments in (**A**) (the circled cell is the one chosen for the image sequences in (**A**)).

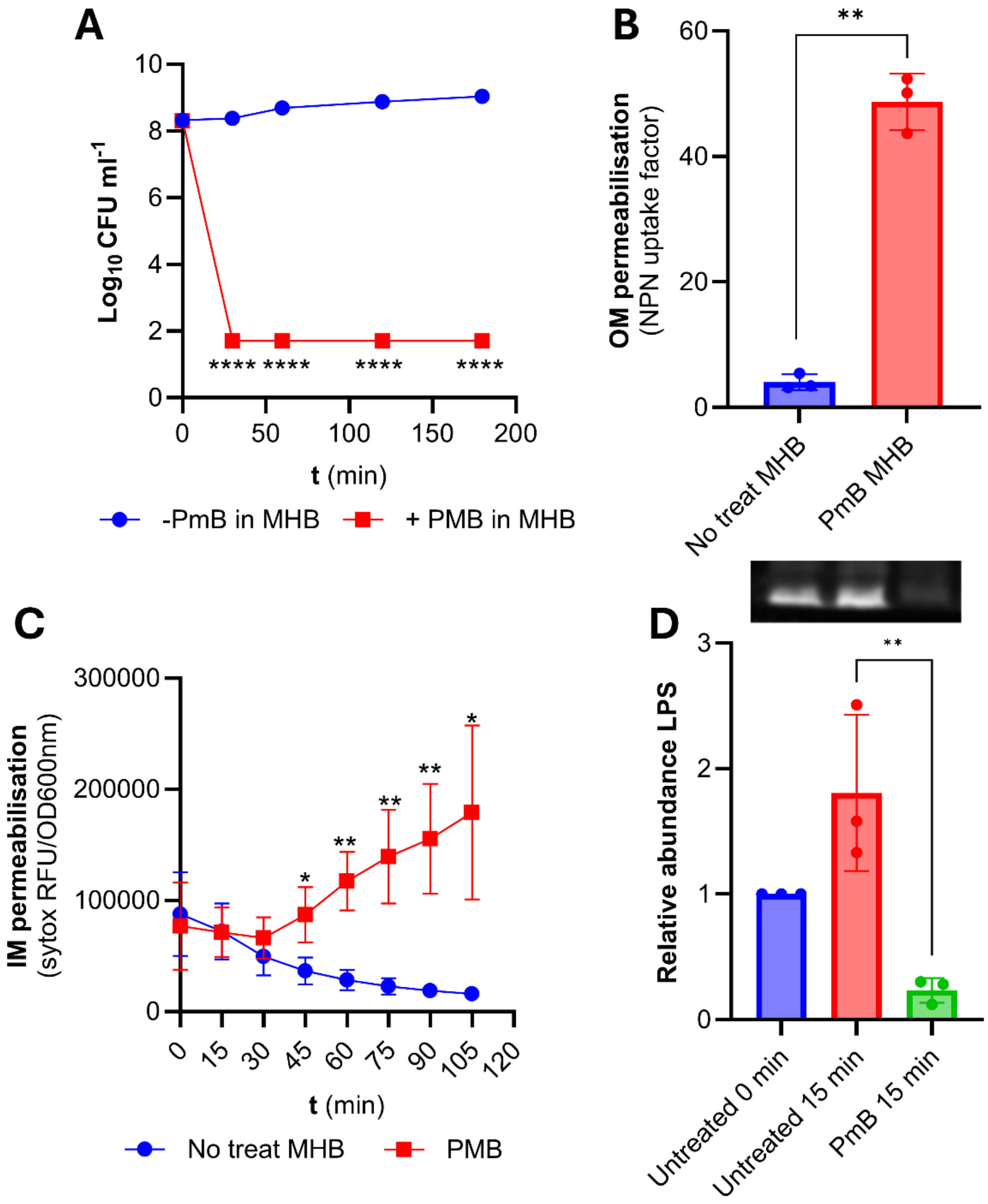


**Supplementary Figure S10. PmB-induced LPS loss occurs in MHB.** To ensure that LPS loss was not an artefact of performing our assays in MM or MM+G, experimentation was repeated in MHB. **(A)** Survival of exponential phase *E. coli* exposed, or not, to 4 µg ml$^{-1}$ PmB or inactive PmB, as determined by CFU counts. **(B)** OM disruption of exponential phase *E. coli* cells during the first 20 min of exposure to 4 µg ml$^{-1}$ PmB in MHB, as determined by uptake of the NPN fluorescent dye. **(C)** OM and IM disruption of exponential phase *E. coli* exposed to 4 µg ml$^{-1}$ PmB in MHB, as determined by uptake of the fluorescent dye SYTOX green. **(D)** Total LPS levels of exponential phase *E. coli* exposed to 4 µg ml$^{-1}$ PmB in MHB for 15 mins. The graph shows the quantification of LPS levels from densitometric analysis using Fiji. For all experiments n=3 independent assays, error bars show the standard deviation of the mean. Significant differences were determined by one-**(B, D)** or two-way **(A, C,)** repeated measures ANOVA. P= *<0.05, **<0.01, ***<0.001, ****<0.0001.

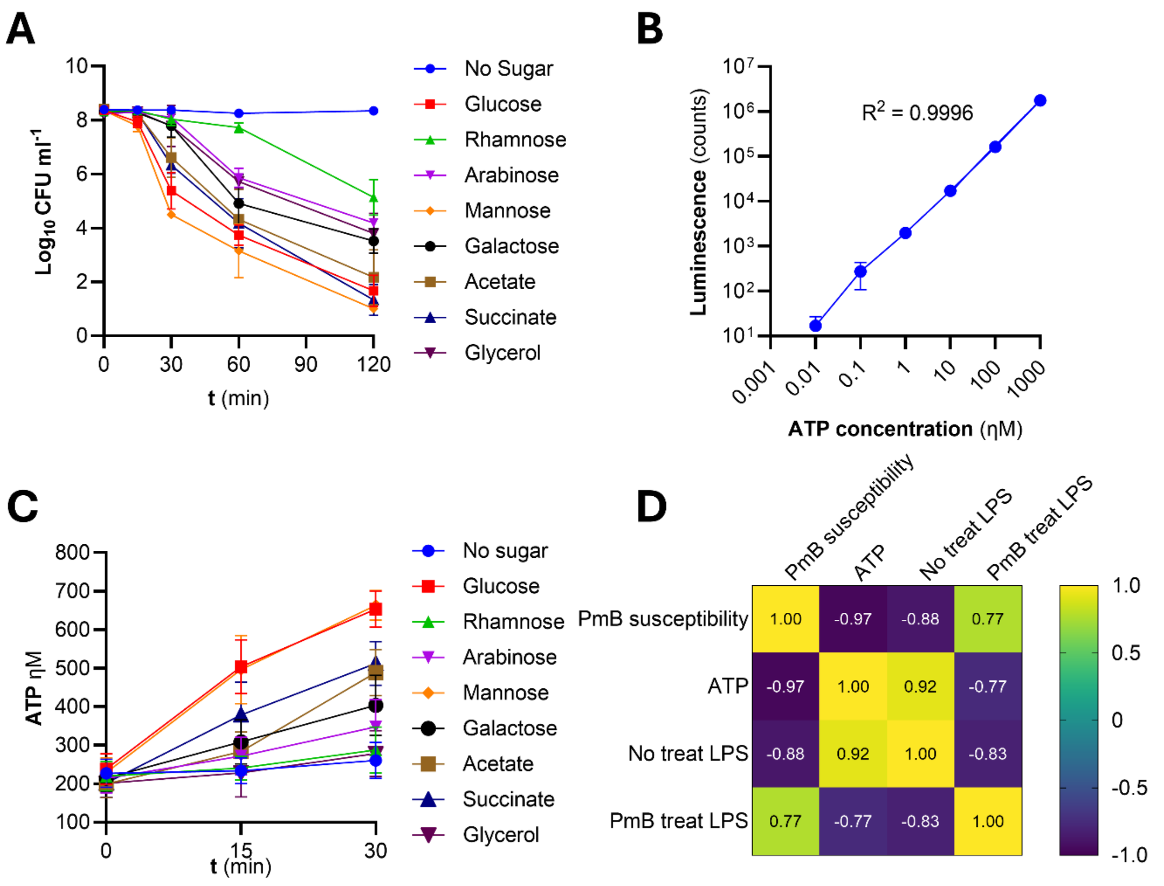


**Supplementary Figure S11. PmB killing requires ATP. (A)** Survival of stationary phase *E. coli* exposed to 4 µg $ml^{-1}$ PmB in MM +/- equimolar concentrations of different sugars, as determined by CFU counts. **(B)** Standard curve plotting ATP concentration (nM) against luminescence counts. Simple linear regression was performed using prism version 10.4.1, where the blue line represents the line of best fit. **(C)** ATP concentration according to standard curve interpolation, of stationary phase *E. coli* incubated in equimolar concentrations of different sugars across a 30-minute time course. **(D)** Four-way correlation matrix showing correlation coefficients of PmB susceptibility, ATP concentration and densitometric analysis pre- and post-PmB treatment. All experiments were replicated in n=3 independent assays. Error bars show the standard deviation of the mean.

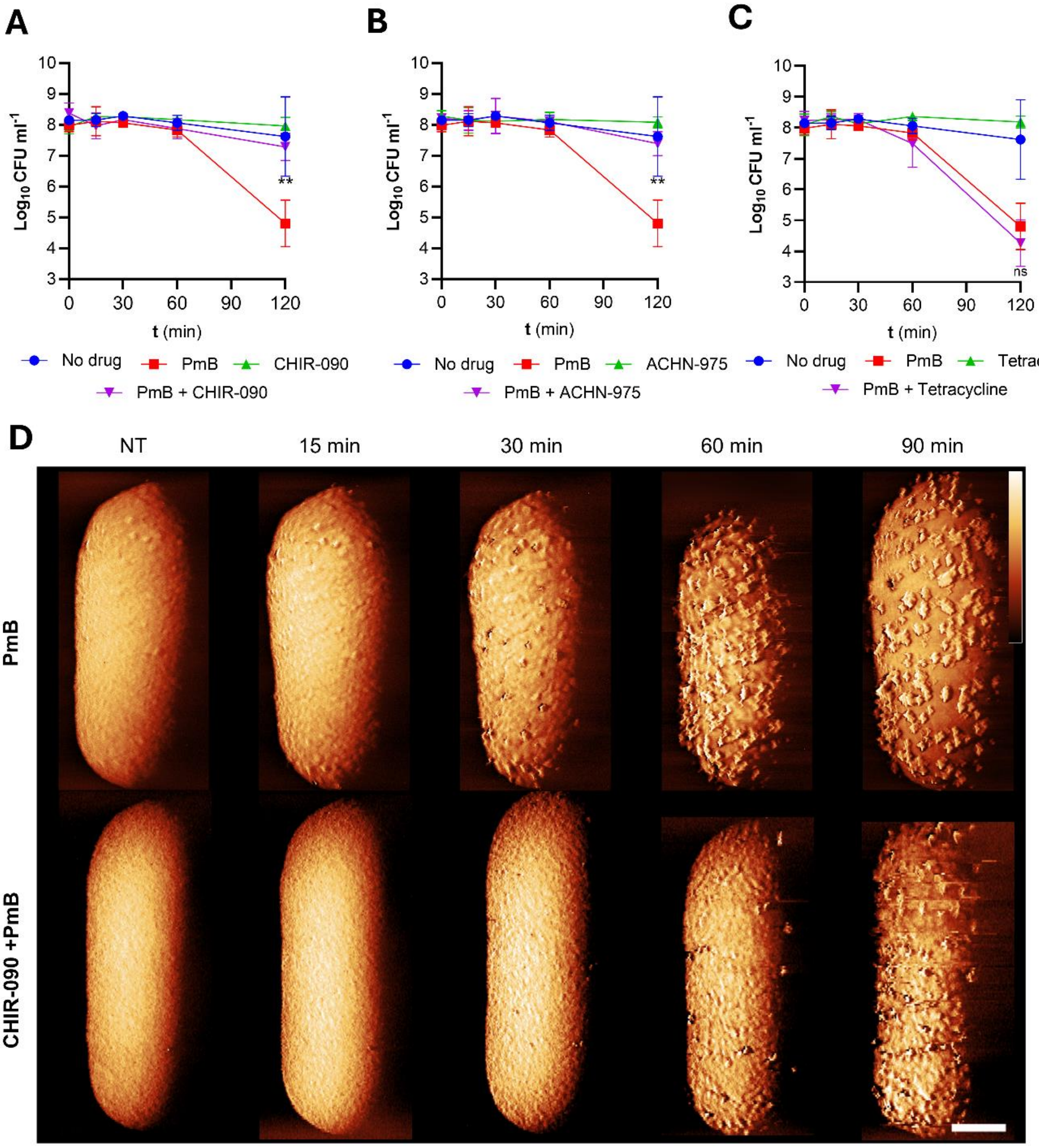


**Supplementary Figure S12. PmB killing requires LPS synthesis. (A, B, C)** Survival of stationary phase *P. aeruginosa* exposed, or not, to 4 µg ml$^{-1}$ PmB in MM+G with or without 1X MIC of LpxC inhibitors CHIR-090 **(A)** or ACHN-975 **(B)** or the protein synthesis inhibitor tetracycline **(C). (D)** AFM phase images showing stationary phase *E. coli* cells exposed to 2.5 µg ml$^{-1}$ PmB in the presence or not of 0.125 µg ml$^{-1}$ CHIR-090 in MM + G, shown as a function of time. Scalebar: 250 nm. Colour scale: 2.5 deg. All experiments were replicated in n=3 independent assays. Error bars show the standard deviation of the mean. Significant differences were determined by two-way repeated measures ANOVA between PmB-treated and PmB + antibiotic-treated conditions. P= **<0.01, ns=not significant.

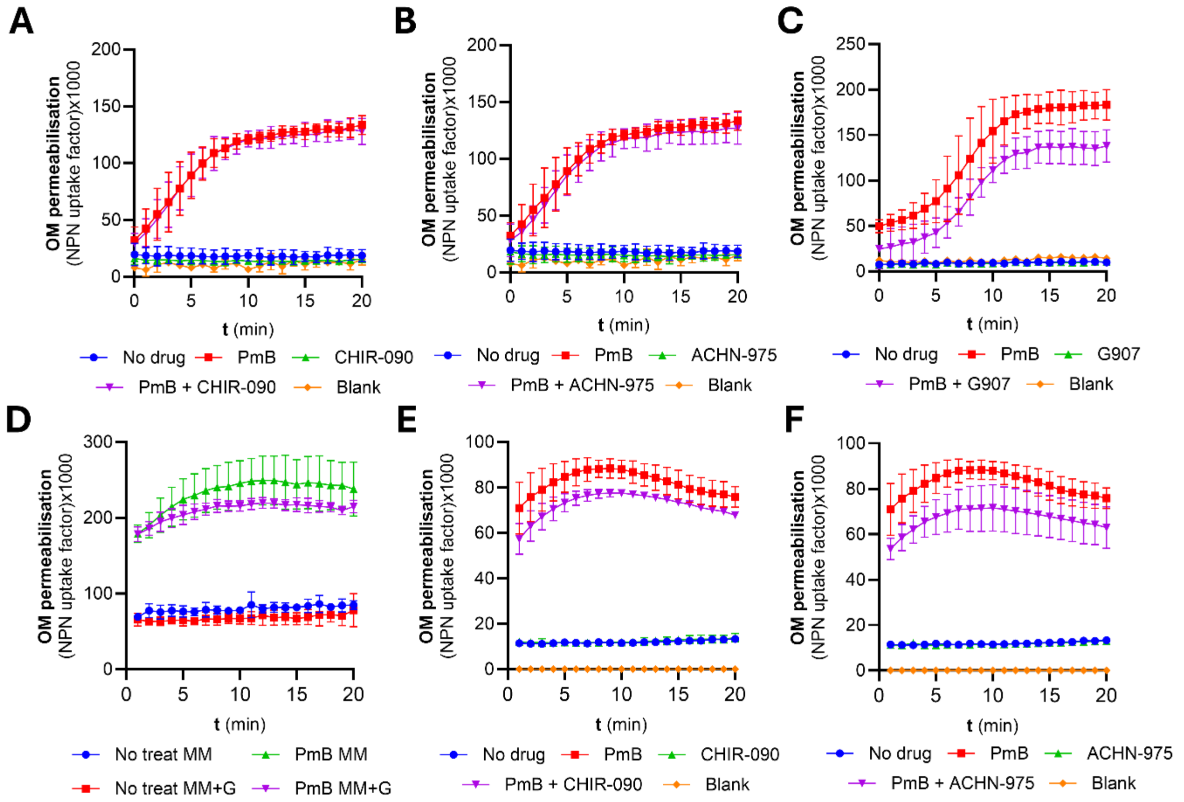


**Supplementary Figure S13. Blocking LPS synthesis or transport does not affect OM permeabilisation. (A, B, C)** OM disruption of stationary phase *E. coli* cells during the first 20 min of exposure to 4 µg ml$^{-1}$ PmB in MM+G with or without 1X MIC of LpxC inhibitors CHIR-090 **(A)**, ACHN-975 **(B)**, or the MsbA inhibitor G907 **(C). (D)** OM disruption of stationary phase *P. aeruginosa* cells during the first 20 min of exposure with and without 4 µg ml$^{-1}$ PmB in MM+G or MM. **(E, F)** OM disruption of stationary phase *P. aeruginosa* cells during the first 20 min of exposure to 4 µg ml$^{-1}$ PmB in MM+G with or without 1X MIC of LpxC inhibitors CHIR-090 **(E)**, ACHN-975 **(F)**. All experiments were replicated in n=3 independent assays. Error bars show the standard deviation of the mean.

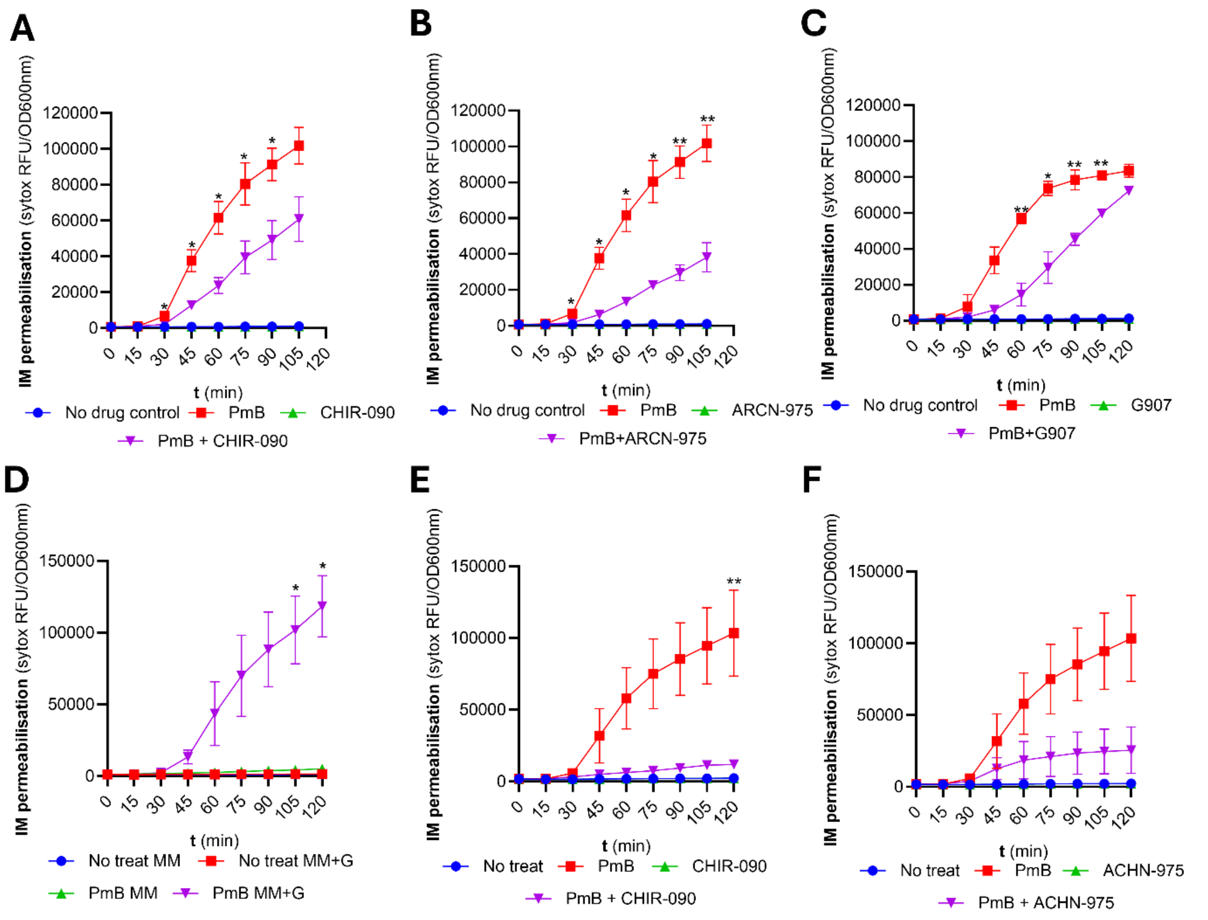


**Supplementary Figure S14. Blocking LPS synthesis and transport reduces IM permeabilisation. (A, B, C)** OM and IM disruption of stationary phase *E. coli* exposed to 4 µg ml$^{-1}$ PmB in MM +G, with or without 1X MIC of LpxC inhibitors CHIR-090 **(A)**, ACHN-975 **(B)**, or the MsbA inhibitor G907 **(C)**, as determined by uptake of the fluorescent dye SYTOX green. **(D)** OM and IM disruption of stationary phase *P. aeruginosa* exposed to 4 µg ml$^{-1}$ PmB in MM +/-G. **(E, F)** OM and IM disruption of stationary phase *P. aeruginosa* exposed to 4 µg ml$^{-1}$ PmB in MM +/-G, with or without 1X MIC of LpxC inhibitors CHIR-090 **(E)**, or ACHN-975 **(F)**. All experiments were replicated in n=3 independent assays. Error bars show the standard deviation of the mean. Significant differences were determined by two-way repeated measures ANOVA between PmB and PmB + inhibitor (**A**, **B**, **C**, **D**, **E**, **F**) or between PmB MM+G and PmB MM (**D**). P= *<0.05, **<0.01, ns=not significant.

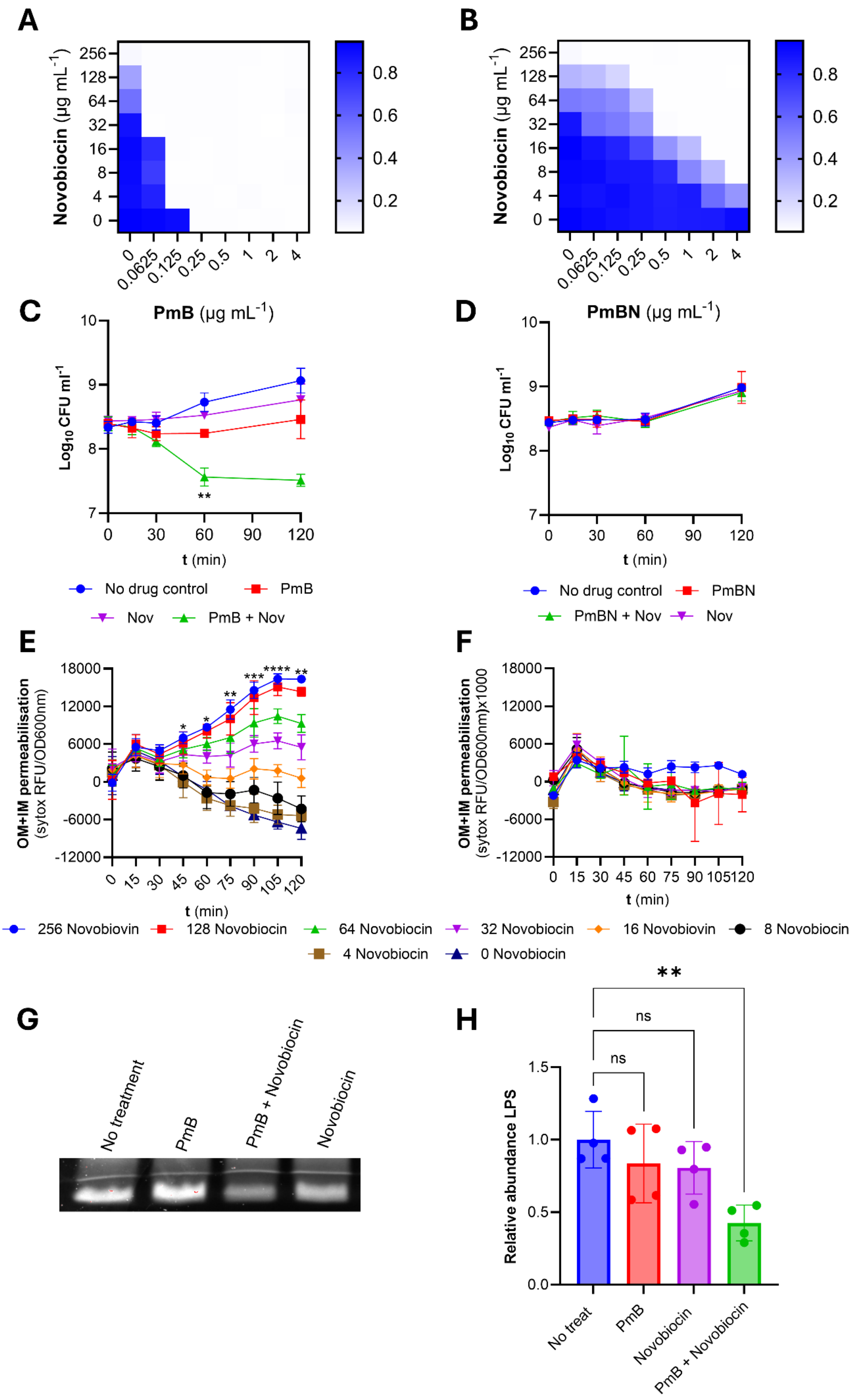
A
Novobiocin (µg mL-1)
256
128
64
32
16
8
4
0
0
0.0625
0.125
0.25
0.5
1
2
4
0.8
0.6
0.4
0.2
B
Novobiocin (µg mL-1)
C
PmB (µg mL-1)
Log10 CFU ml-1
t (min)
No drug control
PmB
Nov
PmB + Nov
D
PmBN (µg mL-1)
Log10 CFU ml-1
t (min)
No drug control
PmBN
PmBN + Nov
Nov
E
OM+IM permeabilisation
(sytox RFU/OD600nm)
t (min)
F
OM+IM permeabilisation
(sytox RFU/OD600nm)x1000
t (min)
256 Novobiovin
128 Novobiocin
64 Novobiocin
32 Novobiocin
16 Novobiovin
8 Novobiocin
4 Novobiocin
0 Novobiocin
G
No treatment
PmB
PmB + Novobiocin
Novobiocin
H
Relative abundance LPS
ns
ns
**
No treat
PmB
Novobiocin
PmB + Novobiocin

**Supplementary Figure S15. Novobiocin promotes PmB-mediated LPS loss and killing.** Previous work has shown that the antibiotic novobiocin promotes the rate of LPS transport from IM to OM via an interaction with LptB [41]. In turn, this leads to increased susceptibility to polymyxin B, although the mechanism was not established [42]. Based on the findings described in this manuscript, we hypothesised that increased LPS transport to the OM would promote PmB-mediated LPS loss, which correlates with bacterial killing by the antibiotic. **(A, B)** Checkerboard broth microdilution assay showing the synergistic growth-inhibitory interaction between novobiocin and PmB **(A),** or PmBN **(B)** against *E. coli*, as determined $OD_{595nm}$ after 18 hr incubation and in line with previous findings [42]. Note: PmBN is an inactive polymyxin analogue that served as a useful control for increased entry of novobiocin caused by OM disruption. **(C, D)** Survival of *E. coli* exposed, or not, to 1 µg $ml^{-1}$ PmB **(C)** or 1 µg $ml^{-1}$ PmBN **(D)** with or without 32 µg $ml^{-1}$ novobiocin. These concentrations were chosen because they showed maximal synergy in the PmBN checkerboard assay. For **(C)**, there was no killing of *E. coli* by the polymyxin or novobiocin alone, but a significant reduction in the viability of *E. coli* in the presence of both antibiotics. By contrast, there was no reduction in bacterial viability in the presence of PmBN with novobiocin. Therefore, bacterial killing in these assays appears to be PmB-mediated, rather than increased ingress of novobiocin into cells caused by OM disruption. **(E, F)** Combined OM and IM disruption of stationary phase *E. coli* exposed to 1 µg $ml^{-1}$ PmB **(E)** or 1 µg $ml^{-1}$ PmBN **(F)** in MHB with and without 32 µg $ml^{-1}$ novobiocin, as determined by uptake of the fluorescent dye SYTOX green. Crucially, in the presence of PmBN, novobiocin did not cause membrane disruption, even at concentrations well above those required to inhibit growth in the checkerboard assay. However, novobiocin promoted PmB-mediated membrane disruption in a dose-responsive manner, confirming that novobiocin promotes the activity of PmB by increasing IM disruption, which is the key step for lethality [27,42]. Next, we wanted to examine the impact of novobiocin on PmB-mediated LPS loss. **(G)** Representative SDS-PAGE image of LPS band intensity of stationary phase *E. coli* exposed, or not, to 1 µg $ml^{-1}$ PmB with and without 32 µg/ml novobiocin for 15 min in MHB. **(H)** Densitometric analysis of the LPS gel in **(G)**, showing that PmB or novobiocin exposure alone at 1 µg $ml^{-1}$ or 32 µg $ml^{-1}$ respectively, had minimal impact on LPS abundance after 15 mins incubation. However, there was a significant drop in LPS levels when PmB and novobiocin were used in combination at these concentrations. Therefore, we concluded that novobiocin enhances PmB activity by promoting LPS loss, most likely via its reported effect on increasing LPS transport from the IM to OM [41,42].

All experiments were replicated in n=3 independent assays. Error bars show the standard deviation of the mean. Significant differences were determined by one- **(H)** or two-way repeated measures ANOVA **(C, D, E, F).** P= *<0.05, **<0.01, ***<0.001, ****<0.0001, ns=not significant.

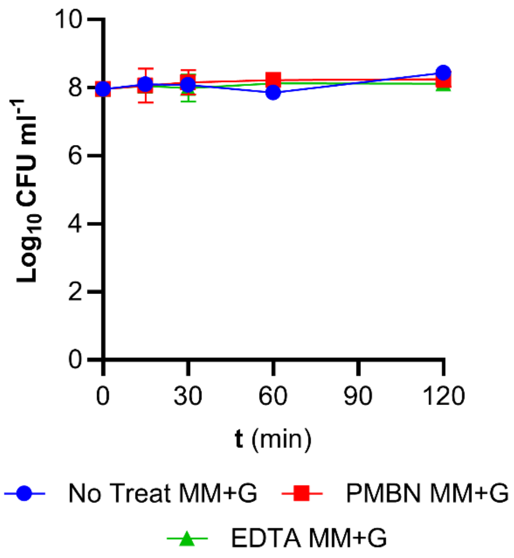


**Supplementary Figure S16. PmBN and EDTA do not have bactericidal activity.** Survival of *E. coli* exposed, or not, to 4 µg $ml^{-1}$ PmBN or 10 mM EDTA in MM+G. All experiments were replicated in n=3 independent assays. Error bars show the standard deviation of the mean.

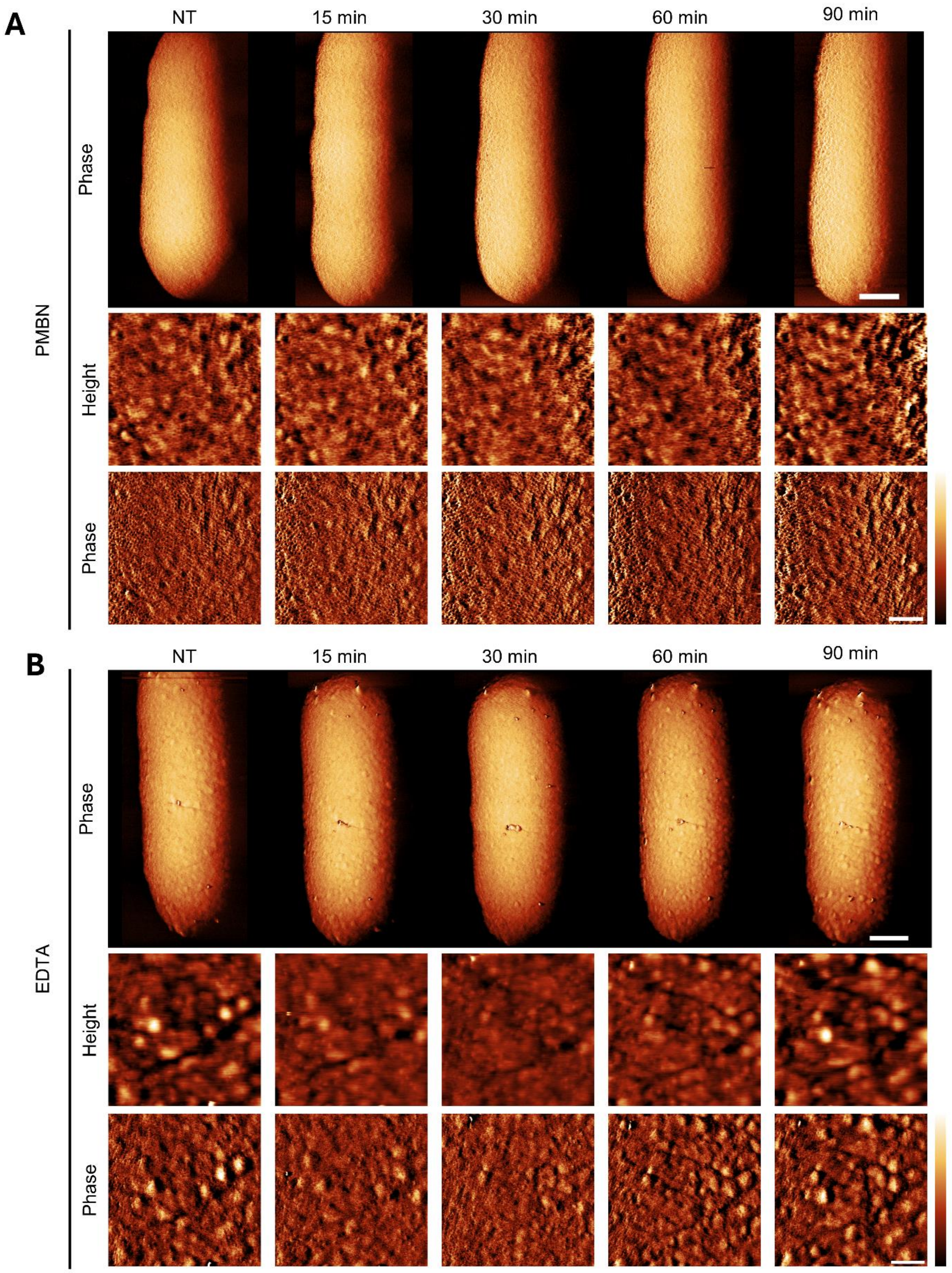


**Supplementary Figure S17. Low-dose PmBN and EDTA exposure has little effect on the OM at the nanoscale**. AFM low and high magnification scans showing stationary phase *E. coli* MG1655 exposed to 2.5 μg ml$^{-1}$ PMBN **(A)** or 10 mM EDTA **(B)** in the presence of glucose for up to 90 minutes. Scalebars: large scans 250 nm, higher-magnification scans 100 nm; colour bar (from top to bottom): **(A)** 4 deg, 3 nm and 0.7 deg, **(B)** 5 deg, 10 nm and 1 deg .

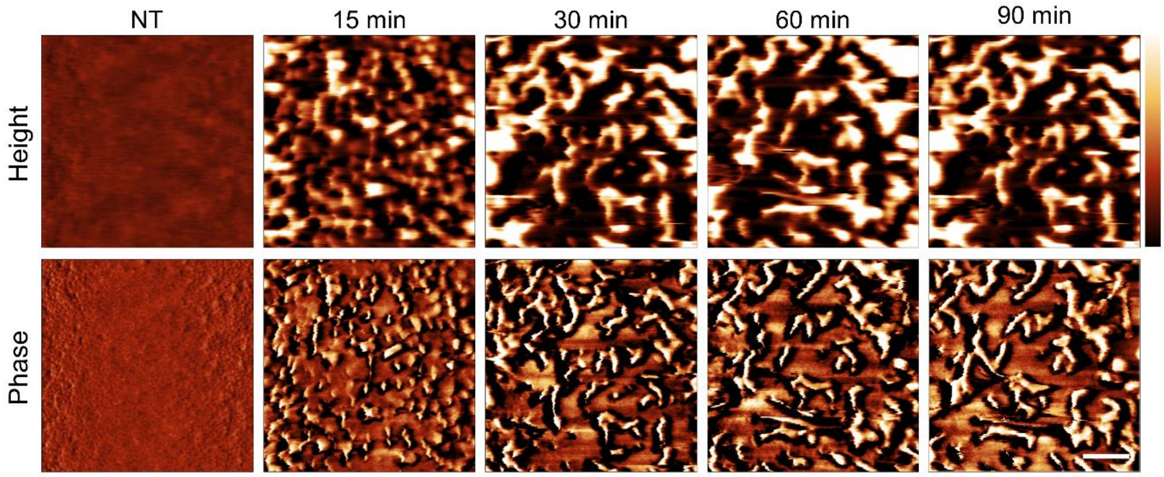


**Supplementary Figure S18. In MM without glucose, co-treatment of stationary phase *E. coli* with EDTA and PmB causes clearly noticeable membrane roughening, resembling the effects observed for PmB in MM + glucose**. **(B)** AFM high magnification scans of the bacteria in Fig. 4H showing stationary phase *E. coli* MG1655 exposed to 10 mM EDTA and 2.5 μg $ml^{-1}$ PmB in M9 salts followed through the time course. Scalebar: 100nm; colour bar: 20 nm (height) and 1.5 deg (phase).
